# Exploring the large-scale properties of a protein secondary structure genotype-to-phenotype map

**DOI:** 10.64898/2026.06.26.734756

**Authors:** Javor K. Novev, Sebastian Schornack, Sebastian E. Ahnert

## Abstract

We perform a large-scale computational characterization of the map of protein primary to secondary structure using an AVR3a class protein effector domain from the plant pathogen *P. palmivora* as a case study. We formulate a modified site-scanning approach for exploring the neutral component of secondary structure phenotypes based on predictions from the machine-learning algorithm Porter 5 and apply it to the AVR3a phenotype. We predict a set of sensitive sites within the effector domain that are generally located at or near the boundaries of structured regions, with restrictions on the possible amino acid residues at these sites dictated by the secondary structure type that they participate in within the WT. We characterize a set of mutated phenotypes derived through the exploration of the neutral component of the WT effector domain, selecting them so that they span a range including both very rarely and very commonly seen secondary structures, and that they include both secondary structures nearly identical to the WT and ones far removed from it. We find that all these diverse phenotypes have an estimated robustness of the same order as that of the WT, and that the robustness scales logarithmically phenotype frequency, as seen in other genotype-to-phenotype maps. Furthermore, we observe that the dependence of the estimated phenotype frequency on the Kolmogorov complexity indicates simplicity bias in the protein secondary structure map.

## 1 Introduction

The mapping of genotypes to phenotypes is a fundamental problem in computational biology that has been the subject of intense study in recent years [1]. Understanding of genotype-to-phenotype (GP) maps is required for a predictive theory of evolution [1], and research has revealed that many fundamental properties of GP maps, such as a high degree of redundancy, i.e., a large number of genotypes mapping onto the same phenotype, and a high robustness, i.e., the property that a high fraction of the possible mutations leave the phenotype unchanged [1, 2]. A GP map that has hardly been studied on a large scale is that of between protein primary and secondary structure [3], i.e., between a protein’s amino acid sequence and its local folding into helical and sheet-like patterns [4]. Here, we study this map’s properties, which have not been studied so far, but are essential to understanding protein evolutionary dynamics, particularly in fast-evolving systems such as viruses and cancer.

We explore the large-scale properties of the protein secondary structure (SS) GP map by studying the neutral space of a chosen model protein, i.e., the region of genotype that maps onto the same phenotype SS as the original protein. We do this via a novel algorithm for efficiently exploring the neutral space known as site-scanning, which for each site of the protein tests whether a randomly chosen mutation is neutral, and mutates it again if it is not, it mutates it again until either it finds a neutral mutation or exhausts all possible point mutations at the site [2, 5]. We identify the protein sites most sensitive to mutations by generating a sample of neutral genotypes through site-scanning, then selecting a random subsample from it and exploring the full point-mutational neighbourhoods of the genotypes from the subsample. We then develop methods to characterize the size and properties of the neutral space of the original phenotype. Furthermore, we probe the evolutionary landscape accessible through mutating the WT protein by analysing the neutral spaces of mutated phenotypes in the same way. For generating secondary structure predictions, we employ Porter 5, an accurate machine-learning-based tool [6].

For our model protein, we choose a variant of the avirulence protein 3a (AVR3a), an effector protein that is essential to the infection of plants with the potato blight pathogen, the oomycete *Phytophthora infestans* [7–9] and other species of the same family. On the one hand, as an effector protein, AVR3a suppresses pathogen-associated molecular pattern (PAMP)-triggered immunity in the plants it infects [7]. Specifically, AVR3a, which contains the conserved RXLR motif [7, 10], suppresses programmed cell-death induced by the *P. infestans* infestin 1 protein [7]. At the same time, effectors such as AVR3a are themselves recognized by plant resistance (R) proteins [11], in this case R3a, that trigger a form of programmed cell death [7].

We focus on a form of the AVR3a effector domain found in *Phytophthora palmivora*, a tropical oomycete that infects perennial crops such as cacao, papaya, mango and oil palm [12]; this form has 74.3% sequence identity to the KI AVR3a variant from *P. infestans* [7] (BLAST E-value: 2.7 × 10^−35^ [13]). Below, we refer to this specific form of the effector domain as the 2A AVR3a variant. Besides shedding light on the general properties of the protein secondary structure map, studying neutral proteins of this mutant, which has to balance efficiency in suppressing cell death with evading immune response, would suggest potential evolutionary trajectories for this economically relevant pathogen.

The methods we develop here are broadly applicable and can be used for, e.g., the study of human pathogens such as the SARS-CoV-2 virus.

## 2 Methods

### 2.1 Secondary structure prediction algorithm

We tested the accuracy of Porter 5 on the NMR solution structure of the AVR3a effector [14]. The secondary structure string predicted by the fast version of the algorithm for 72.5% of the sites agreed with the one parsed from [14] via DSSP [15] version 3.1.4. The prediction of the more accurate version of Porter 5 was in agreement with the NMR solution structure for 75.4% of the sites. We also generated predictions with AlphaFold 2 [16] in its parallel implementation Parafold [17], which were between 71.5% and 74.5% accurate by the same measure. However, both Porter 5 and AlphaFold 2 were trained on data from the PDB and the 2NAR structure is in the training sets for AlphaFold 2 and likely Porter 5 as well, likely making it a poor indicator of their accuracy in predicting the secondary structure of sequences not found in the PDB such as the ones we study here. Considering the computational cost of folding hundreds of thousands of mutants and the accuracy of all these algorithms on a system closely related to our WT protein, we chose to use the fast version of Porter 5 for our large-scale study.

### 2.2 Site-scanning for exploring protein secondary structure neutral components

We find neutral mutants through a modified version of the site-scanning approach developed in [2] and modified in [5]. Starting with the wild type (WT) sequence, we mutate sites one by one, choosing the mutated site index according to a random permutation. We substitute the amino acid residue at the chosen site with a randomly chosen one that is accessible via a point mutation in DNA, then use Porter 5 [6] to predict its secondary structure.

If the mutation is neutral, i.e., if it has the same predicted secondary structure as the WT, we retain it and move on to mutating the next mutation chosen according to the random permutation. We repeat this process until we have generated the desired number of neutral mutations (200); we perform this process 10 times with different randomizations in parallel starting from the WT sequence.

We then select a subsample of the sample of neutral mutants (50 for the WT; 20 for the other phenotypes - see section 3.1) and predict the secondary structure of all mutants in its point mutational neighborhood, i.e., all sequences accessible via a point mutation in DNA.

### 2.3 Data analysis

Within each of the 10 samples, we calculate the fraction of point mutants to each of these genotypes from in the subsample that are neutral. For each sample ‘i’, genotype ‘j’ and site ‘k’ this fraction is equivalent to a site-specific genotypic robustness:

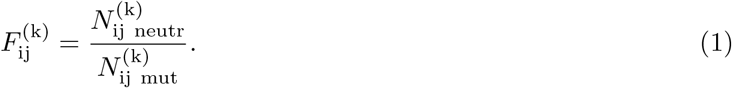

Note that the number of mutants accessible via a point mutation in DNA, 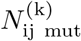, depends on both the starting sequence ‘j’ and the site ‘k’. The robustness of genotype ‘j’ is the average of the neutral mutant fraction 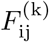 over all *L* sites in the protein [1]:

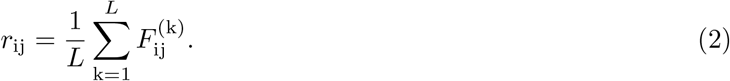

We calculate an estimate of the phenotypic robustness for the sample 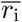 as the average of *r*_ij_ over all *N*_subsample_ genotypes in the subsample:

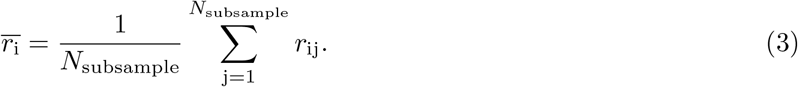

We then estimate the phenotype robustness as an average of 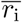 over all samples:

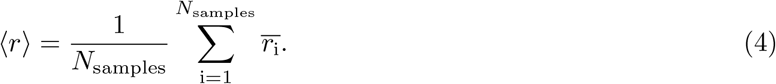

For each sample, we compute the standard deviation 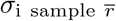 associated with 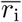:

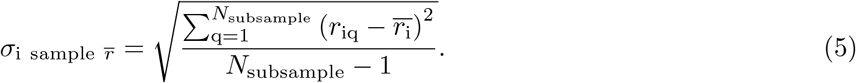

We then pool the sample standard deviations as follows [18]:

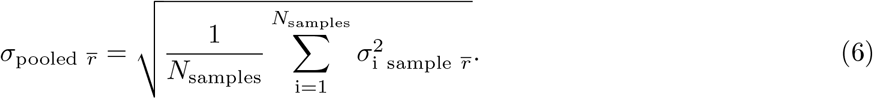

We then calculate the standard error of the mean as

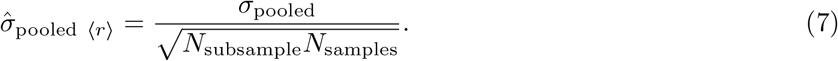

By analogy, we define the overall average of the fraction of neutral mutations for each site, ⟨*F*^(k)^⟩ and its pooled standard error of the mean, 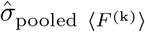.

We additionally measure the average number of changes to the secondary structure that a non-neutral point mutation induces. We calculate this quantity for each site (indexed ‘k’) and normalize it to the sequence length,

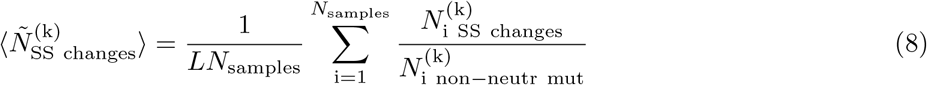

We calculate 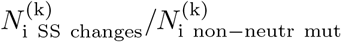 for entire samples rather than individual genotypes because many genotypes have sites at which all point mutations are neutral - for them, the equivalent measure at the genotype level is undefined. For the same reason, we calculate the standard deviation of 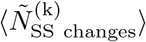 as

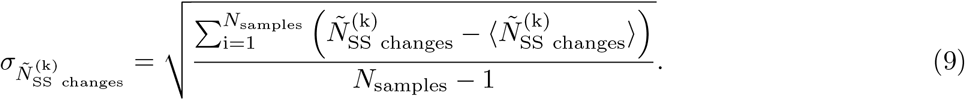

We then use the associated standard error of the mean, 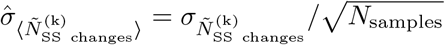.

We estimate the the size of the neutral component based on the fraction of neutral mutations averaged over all subsample genotypes in all samples:

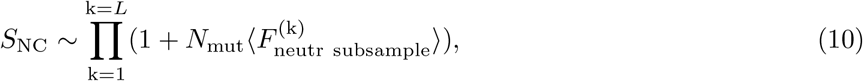

where *N*_mut_ = *S*_alphabet_ −1 is the number of point mutations possible at the site, where *S*_alphabet_ = 20 is the number of amino acid residue types in naturally occurring proteins.

We then use the neutral component size to estimate the phenotype’s frequency *f*, i.e., the fraction of all possible genotypes that map to it [1]:

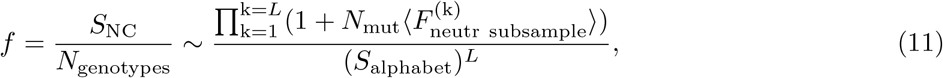

where we have assumed that only the single neutral component that we have characterized makes up a significant part of the neutral set for the phenotype under study. A general property of many GP maps is that ⟨*r*⟩ scales with the logarithm of the phenotype frequency *f*, or, equivalently, of the neutral set size [1, 19].

While exploring the neutral component of the 2A WT secondary structure phenotype, we generate 311393 mutated sequences, ~ 45.8% of them having the WT phenotype. We select 14 mutant phenotypes whose frequency in this set spans several orders of magnitude, ranging from 0.02% (52 occurrences) to 1.8% (5614 occurrences); they rank between the 4th most frequent and 244th most frequent phenotype in the set - see Table 1. We choose to characterize the neutral components of these mutated phenotypes because they span a wide range of distances from the WT - in terms of secondary structure, their Hamming distance from the WT ranges between 1.3% and 75.6% - and they are seen at appreciable frequencies, which makes them less likely to be purely artefacts of the secondary structure prediction algorithm. For the phenotypes ranked 6, 24 and 244, we choose a ‘WT’ sequence that to study via site-scanning at random from all in the 2A dataset that map onto the respective phenotype. For all other mutant phenotypes, we analyze the distribution of Hamming distances between the sequences (primary structures) mapping onto the mutated phenotype and the sequence of the 2A AVR3a variant. We then pick a ‘WT’ sequence for the mutated phenotype at random from those whose Hamming distance is equal to the median of the distribution.

**Table 1.**
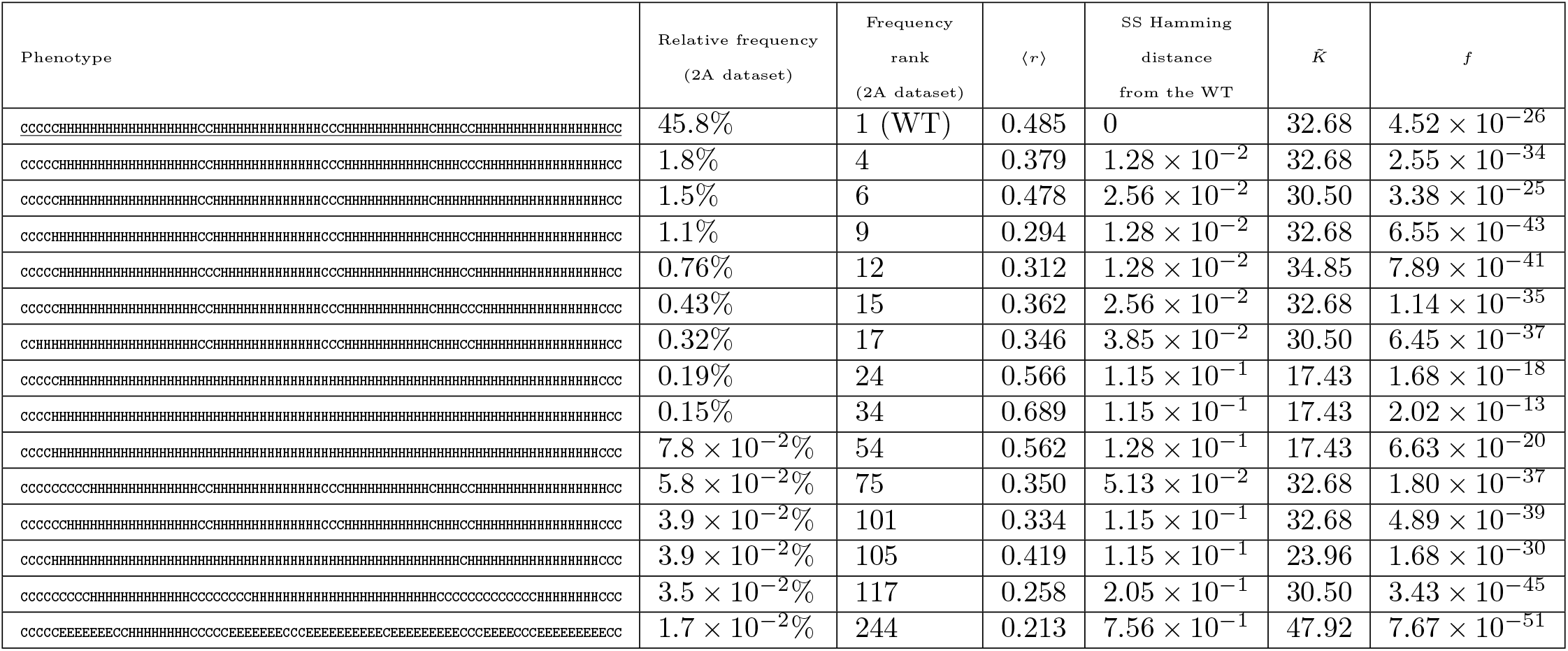
Properties of some secondary structure phenotypes encountered through exploring the neutral component of the 2A variant of the AVR3a effector protein. We select the mutated phenotypes listed here for further study as they span a wide range of relative frequencies within the set of 2A-derived mutants, secondary structure Hamming distances from the WT, and approximate Kolmogorov complexities 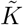. We study their neutral components using the same approach we applied to 2A, which yields robustnesses ⟨*r*⟩ of the same order of magnitude as the WT despite the large differences in frequency within the 2A dataset. The estimated global frequencies *f*, however, differ by many orders of magnitude between these phenotypes, see also Fig. S6 in the SI.

### 2.4 Tertiary structure predictions and analysis

We calculate TM-align scores [20] to compare the similarity of AlphaFold-predicted structures for AVR3a 2A-derived mutants to the wild type. We check whether groups of mutants have the same TM-align score distributions via the one-sided Wilcoxon rank-sum test and calculate the associated *p*-values. We parse the secondary structures for all mutants studied with AlphaFold 2 via DSSP [15] and calculate the Hamming distance between their secondary structure strings and that of the WT. We then compare distributions of secondary-structure Hamming distances in the same way as those of TM-align scores.

## 3 Results

### 3.1 Optimizing the subsample size and type

We study the point mutational neighborhoods of a subsample of 50 randomly chosen neutral genotypes within each of the 10 samples of 200 neutral genotypes generated from the 2A AVR3a effector domain sequence. We calculate estimates for the phenotype robustness ⟨*r*⟩ as described in the Methods based on this data, then calculate new estimates based on sparser subsamples chosen from the original subsamples of size 50. We study sparse subsamples ranging in size from 5 to 25 genotypes, choosing which genotypes to include in three different ways: 1) randomly; 2) aiming for them to be equidistant in sequence from the WT and 3) aiming for them to be equidistant from each other in terms of the number of site-scanning steps required for generating them from the WT. We find that for the WT phenotype, randomly chosen sparse subsamples consistently yield ⟨*r*⟩ estimates intermediate between those for the other two subsample types, and that for each subsampling strategy the results approximately converge with respect to subsample size when 25 genotypes are used, as seen in Figure 1. For the other phenotypes, we observe that ⟨*r*⟩ has already converged for subsamples of 20 neutral genotypes (see Supplementary Figure S4 for an example) and calculate robustness estimates from randomly selected subsamples of that size. We show a visualization of all explored neutral mutants derived from the 2A AVR3a sequence in Figure S2, where sequence communities determined via the Louvain algorithm are plotted in the same colour.

**Figure 1.**
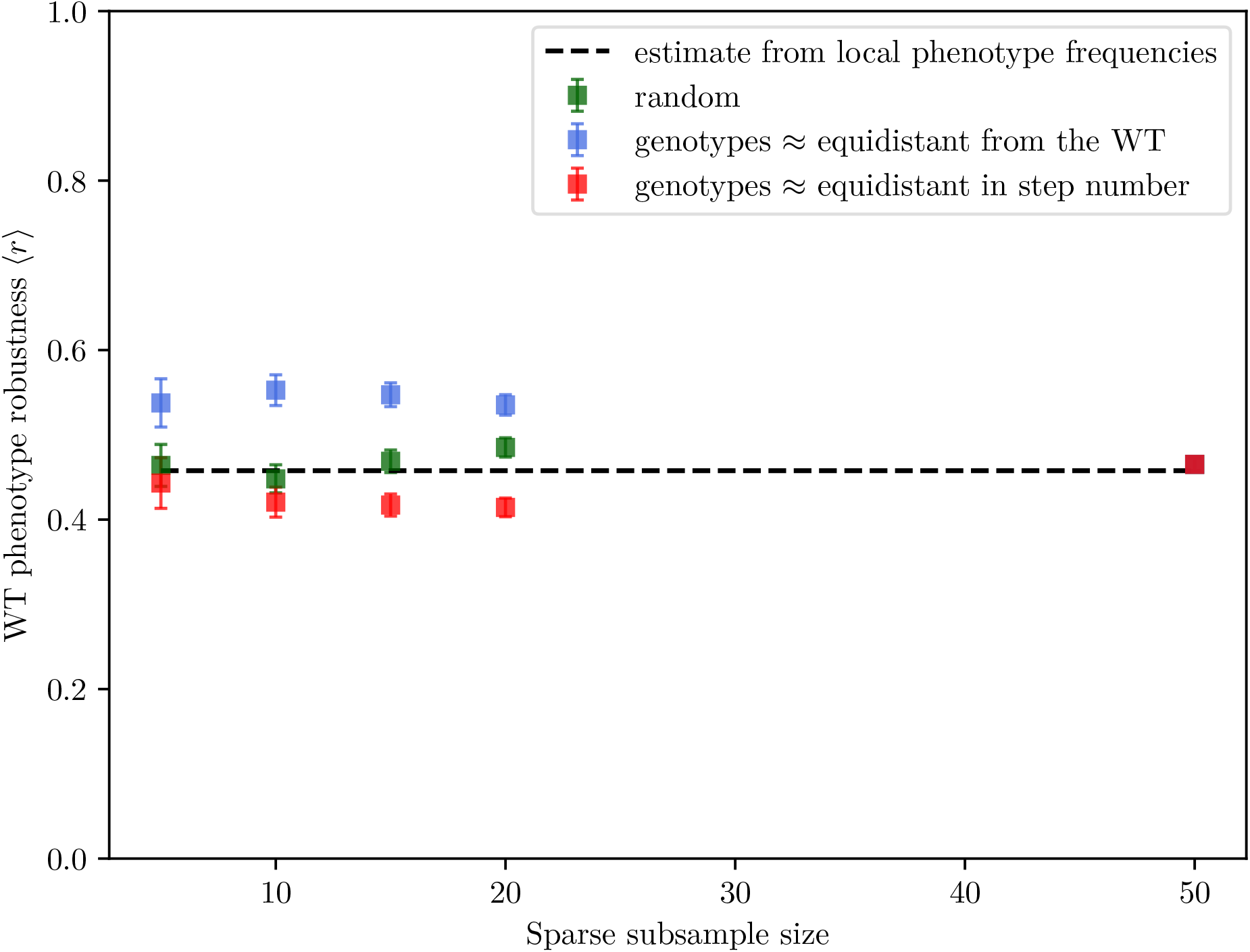
The estimate for the robustness of the 2A AVR3a secondary structure approximately converges at sparse subsample size ~25. Using a random subsample yields robustness intermediate between that estimated by choosing a subsample of neutral genotypes that are approximately equidistant from the WT sequence and that from subsamples of genotypes that are approximately equidistant in the number of site-scanning steps required to derive them from the WT. The dashed line shows the estimate based on the fraction of genotypes encountered in the site-scanning run that map onto the WT phenotype.

### 3.2 Sites identified as sensitive in the 2A WT phenotype

As seen in Figure 2, the most sensitive site according to our algorithm is site 59 with an average fraction of neutral mutations ⟨*F*^(59)^ ⟩ = 2 × 10^−3^. Moreover, all but 6 × 10^−3^% of all neutral mutants of the 2A AVR3a phenotype have glycine at this site. The reason is that in the prediction for the WT 2A phenotype 58 and 59 are the only sites separating two *α*-helices. Glycine is known to destabilize *α*-helices due to its conformational flexibility [21, 22] and to have a preference for occurring as a helix-terminator that is strongly conserved in homologous sequences [23]. Thus, mutating site 59 to a different amino acid generally increases the probability that the *α*-helix extends to this site. Indeed, the secondary structure at site 59 is predicted to be helical in ≈6.1% of the mutated sequences generated during our exploration of the neutral component of the 2A phenotype, in ≈6.8% the same is true of site 58, and in ≈4.3% both sites are predicted to be part of an *α*-helix. As we see when exploring the neutral components of mutated phenotypes, many of those with an extended helix in that region have a robustness similar to that of the WT or higher, see Table 1.

**Figure 2.**
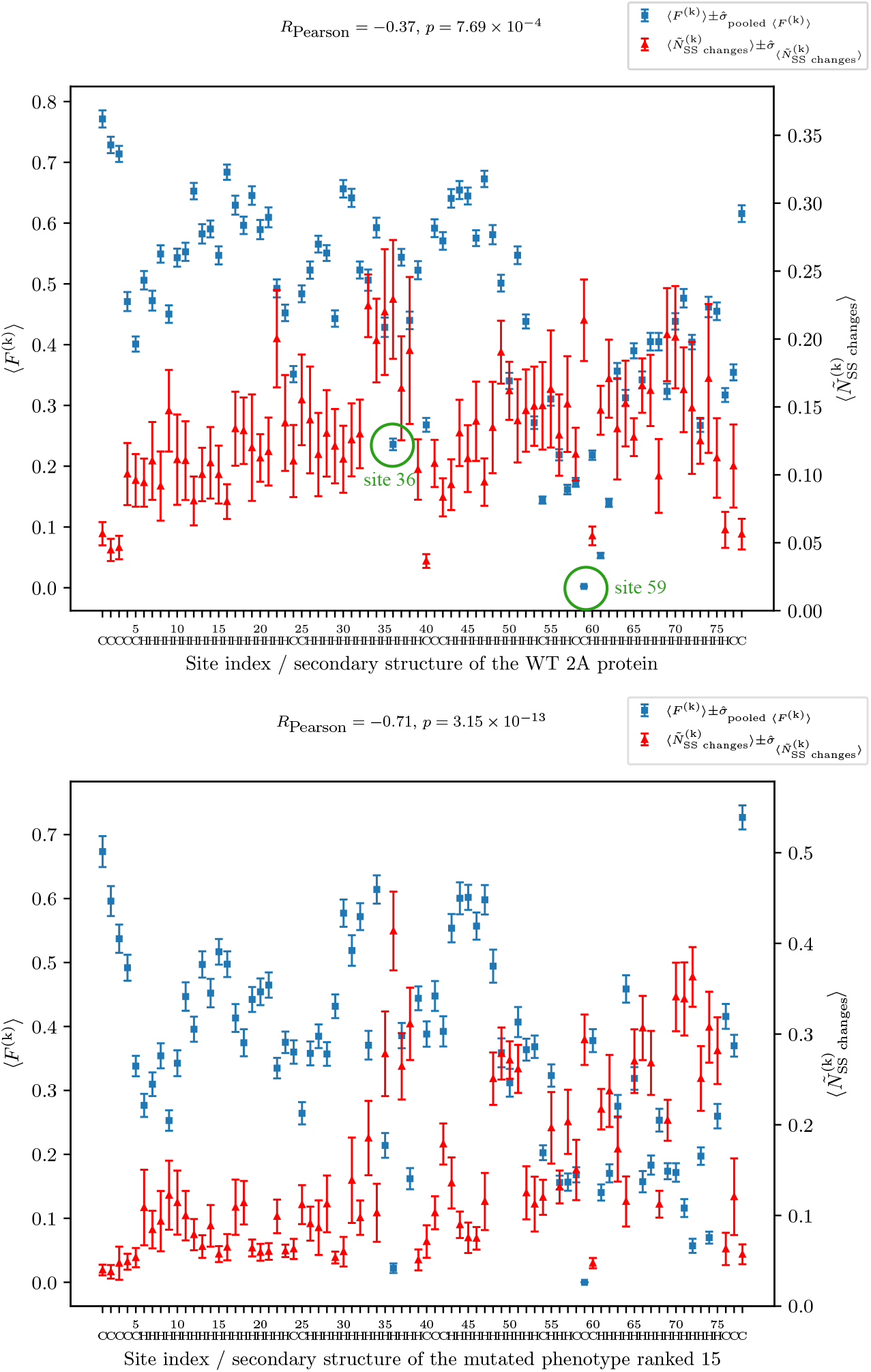
The fraction of neutral mutations (blue squares) averaged over all subsamples and samples ⟨*F*^(k)^⟩ measures the sensitivity of sites within the 2A AVR3a effector domain. Low values of ⟨*F*^(k)^⟩ point to sites where mutations are particularly likely to disrupt the secondary structure of the effector domain. ⟨*F*^(k)^⟩ is generally anticorrelated with the normalized average number of changes induced by point mutations at a given site 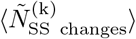. *Top*: 2A AVR3a WT phenotype, *bottom* : phenotype ranked 15 in the 2A dataset. Mutating sites 36 and 59 in genotypes with the phenotype of the 2A AVR3a variant (see main text) affect particularly large numbers of other sites, as also seen in Suppl. Figure S1. The site that allows the least variation in genotypes with the 2A AVR3a phenotype is 59 and is occupied by glycine (G) in virtually all neutral mutants. Glycine tends to destabilize helices and its presence at this position, which is predicted to be at the boundary between a structured and an unstructured region, is likely key for maintaining the secondary structure of 2A AVR3a. Most other highly 7sensitive sites are also found at or near the boundaries between secondary structure elements (e.g., sites 4-5, 24, 40, 58-63 for the WT phenotype) as also seen in the panels for mutated phenotypes. Site 36 stands out as highly sensitive without following this pattern but it is known that tryptophan (W) there together with tyrosine (Y) in the WT forms part of a conserved WY motif. Hydrophobic interactions between them maintain the tertiary structure of AVR3a-class proteins, see the main text and Figure 3.

Preserving the 2A phenotype at other highly constrained sites, also tends to require that the mutated residue preferentially form the same secondary structure element as the residue in the WT. Approximately 75% of the studied sequences with the 2A phenotype have lysine at the second most sensitive site in the sequence (⟨*F*^(61)^⟩ = 5 × 10^−2^), which is predicted to be within a helix. Lysine preferentially forms *α*-helices according to the analysis in [24], and so do most of the other residues found at that site within the neutral component, adding up to ≈90% ‘helix-preferrers’ at the site. Similarly, at the fourth most sensitive site (⟨*F*^(54)^⟩ = 1.3 × 10^−1^), is predicted to be the only coil residue separating two helical regions and has a serine residue in the WT. Like glycine, serine has a tendency to destabilize helices [22, 24]; ≈61% of neutral mutants have serine at this site, and ≈92% of them have a residue that does not preferentially form *α*-helices. While site 59, 61 and 54 are some of the most striking examples, they fall within a general pattern of local minima in the average neutral mutation fraction ⟨*F*^(k)^ ⟩ observed at sites on or adjacent to the boundaries of structured regions (e.g., sites 4-5, 24, 40, 58-63, Figure 2). This pattern is also seen in our data on mutated phenotypes, for example for the ones ranked 6, 34 and 54 in the AVR3a 2A dataset (see Table 1), which are represented in Figure 2. The phenotype with rank 34 has the longest continuous helix and displays this pattern particularly clearly. It should be borne in mind that Porter 5 predicts *five* separate *α*-helices for the 2A AVR3a variant. It is difficult to establish what the number of helices is as crystallographic data on related proteins indicates a fold with *four* helices [25], whereas NMR solution data gives *three* [14]; this may in part reflect the different conditions used for obtaining these structures. Besides, it is possible that the fold of the 2A variant from *P. palmivora* is in fact modified with respect to that of *P. infestans* AVR3a, and the predicted short breaks in the otherwise helical structure at sites 54 and 59 are likely accurate because the amino acid residues in the 2A sequence at these sites have a strong tendency against forming *α*-helices.

AVR3a and related effectors exhibit an evolutionarily conserved WLYY motif - the amino acid residues tryptophan (W), leucine (L) and tyrosine (Y) at specific sites are attracted through hydrophobic interactions that stabilize the *α*-helix conserved protein fold [25]. Alignment of the AVR3a sequence in [25] to the 2A AVR3a effector variant that we study reveals that the residues participating in the WLYY motif are as follows: W at site 36, L at site 50, Y at sites 66 and 69 within the 2A AVR3a effector domain [25]. Among these sites, 36 and 50 are highlighted as relatively sensitive by our site-scanning study. As we study only the secondary structure of mutated proteins, it is not surprising that the other sites are not identified by our method since they affect protein organization on a larger scale. As we exclusively focus on the secondary structure of mutants, we also disregard instances where there the function of the effector protein may be require specific amino acid residues at specific sites. We perform two limited small-scale studies of the tertiary structure of mutants with up to 4 amino acid substitutions in the following sections.

### 3.3 The number of changes to the secondary structure correlate with site sensitivity

Figure 2 shows that the normalized average number of changes that a non-neutral point mutation at a given site induces in the secondary structure (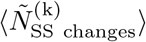, red triangles) is generally anticorrelated with the fraction of neutral mutations at the site (⟨*F*^(k)^ ⟩, blue squares). This means that mutations at the *most* sensitive sites (most notably 36 and 59 in the 2A AVR3a phenotype) also tend to lead to phenotypes that are most distant from the WT. The same general statistically significant trend is observed not just for the 2A AVR3a phenotype, with a typical Pearson correlation coefficient in the range ~ (−0.7, −0.3) - see Figure S3 in the SI for details. The exceptions are the genotypes ranked 34 and 54, for which the correlation is not significant, and 105, for which *R*_Pearson_ *>* 0; all of these contain longer continuous helical regions than the WT.

The anticorrelation between ⟨*F*^(k)^⟩ and 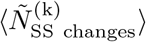 seen in most phenotypes is in contrast with the ‘phenotypic cliffs’ observed in the RNA secondary structure genotype-to-phenotype map [26], i.e., with the tendency for mutations at the *least* sensitive sites to cause the largest phenotypic changes to the minimum free energy (MFE) structure. These differences likely come from the fundamental properties of the two genotype-to-phenotype maps. In RNA secondary structure, changing a robust site, which tends to be unpaired, may easily have a long-range effect on the MFE structure if it is energetically favourable for the mutated base to pair with a distant base. Point-mutation induced switching between protein folds has been observed [10, 27], but is generally rare [28]. As the point mutations affect protein folding via subtle effects such as changing solvent accessibility [10] and hydrophobic interactions [27] rather than relatively strong ones such as the hydrogen-bond-based complementary interactions in RNA, it is not surprising that their effect on secondary structure tends to be local. The sites where non-neutral point mutations induce the largest secondary structure changes in our dataset also tend to be the most sequence-restricted ones, likely because changing the residue at such a site switches the secondary structure of its neighbours. A clear example of that is site 59, where mutating the helix-breaking glycine tends to extend the neighbouring helix.

### 3.4 Testing sensitive sites identified via Porter 5 with AlphaFold 2

We further investigate several sites via AlphaFold 2 [16]. We generate all point mutants for three sites identified as sensitive with respect to secondary structure by our site-scanning algorithm: 36, 59 and 61, as well as site 1, which our method identifies as highly robust (see Figure 2). We compare the AlphaFold 2 predictions for the secondary and tertiary structure of these mutants to a set of 20 others in which the amino acid a random site (except 1, 36, 59 and 61) is mutated to any of the possible 19 alternative residues. We find that mutations at site 36 are predicted to be significantly less similar to the WT than random ones in terms of both secondary structure Hamming distance from the WT (*p* : 5.72 × 10^−3^) and TM-align score (*p* : 1.81 × 10^−5^). Similarly, mutations at site 59 introduce greater changes to both secondary (*p* : 2.46 × 10^−2^) and tertiary structure (*p* : 9.08 × 10^−4^) than ones at a random site. This is in line with the observation from Figure 2 that mutations at these sites cause large changes in secondary structure. AlphaFold 2 predictions show that mutations at site 1 cause changes in secondary and tertiary structure that are not significantly different from those induced by a mutations at a random site, in agreement with the conclusion from our Porter 5 study that site 1 is the most robust one in the sequence. In contrast, sequences with mutations at site 61 do not differ significantly from ones with a randomly mutated site in terms of either their secondary structure Hamming distance or their TM-align score against the WT. We show the distributions of TM-align scores and normalized Hamming distances for these mutants in Supplementary Figure S7. Experimental testing would be the best way to establish the true sensitivity of this site.

### 3.5 AlphaFold 2 predictions confirm the mutational sensitivity of sites involved in the WLYY motif

We perform a small-scale study of the sensitivity of the sites involved in the WLYY in terms of protein tertiary structure. We use AlphaFold 2 [16] to predict the folding of sets of sequences that contain between 1 and 4 point substitution mutations at sites 36, 50, 66 and 69. For every number of point mutations studied, we construct a set of 10 sequences in which the hydrophobic WLYY residues are changed to other residues classified as hydrophobic in [29] and another set in which they are replaced with charged residues. We construct a control set of 10 mutants with 1 to 4 randomly chosen changes in which residues at all sites except 36, 50, 66 and 69 can be mutated to any of the 19 possible alternatives.

We find that mutants in which 2 to 4 of the residues at sites from the WLYY motif are replaced with other hydrophobic ones have a significantly lower TM-align score than mutants with the same number of changes at random other sites - *p*: 2.06 × 10^−2^, 2.47 × 10^−2^, 1.25 × 10^−3^ for double, triple and quadruple mutants, respectively. In contrast, the distributions of secondary-structure Hamming distances are not significantly different between the WLYY and the random mutants. WLYY mutants with charged residues at these sites have TM-align score distributions that differ more significantly from those of random mutants - *p*: 1.42 × 10^−2^, 1.60 × 10^−3^, 5.76 × 10^−4^, 4.40 × 10^−4^ for 1, 2, 3 and 4 changes. Only the secondary structure of mutants in which three of the WLYY residues are substituted with charged hydrophilic ones differs significantly from that of random mutants with the same number of changes (*p* = 4.45 × 10^−2^). We illustrate several of the most striking cases in Figure 3, which demonstrates that double mutants with hydrophobic amino acids at sites 36 and 69 have a nearly identical fold to the wild type, particularly when the W and Y residues are swapped (TM-align score: 0.98), whereas a mutant with two positively charged residues at these sites markedly diverges from the tertiary structure of 2A AVR3a. We show a more detailed comparison with the residues drawn explicitly in Figure S8 in the SI.

**Figure 3.**
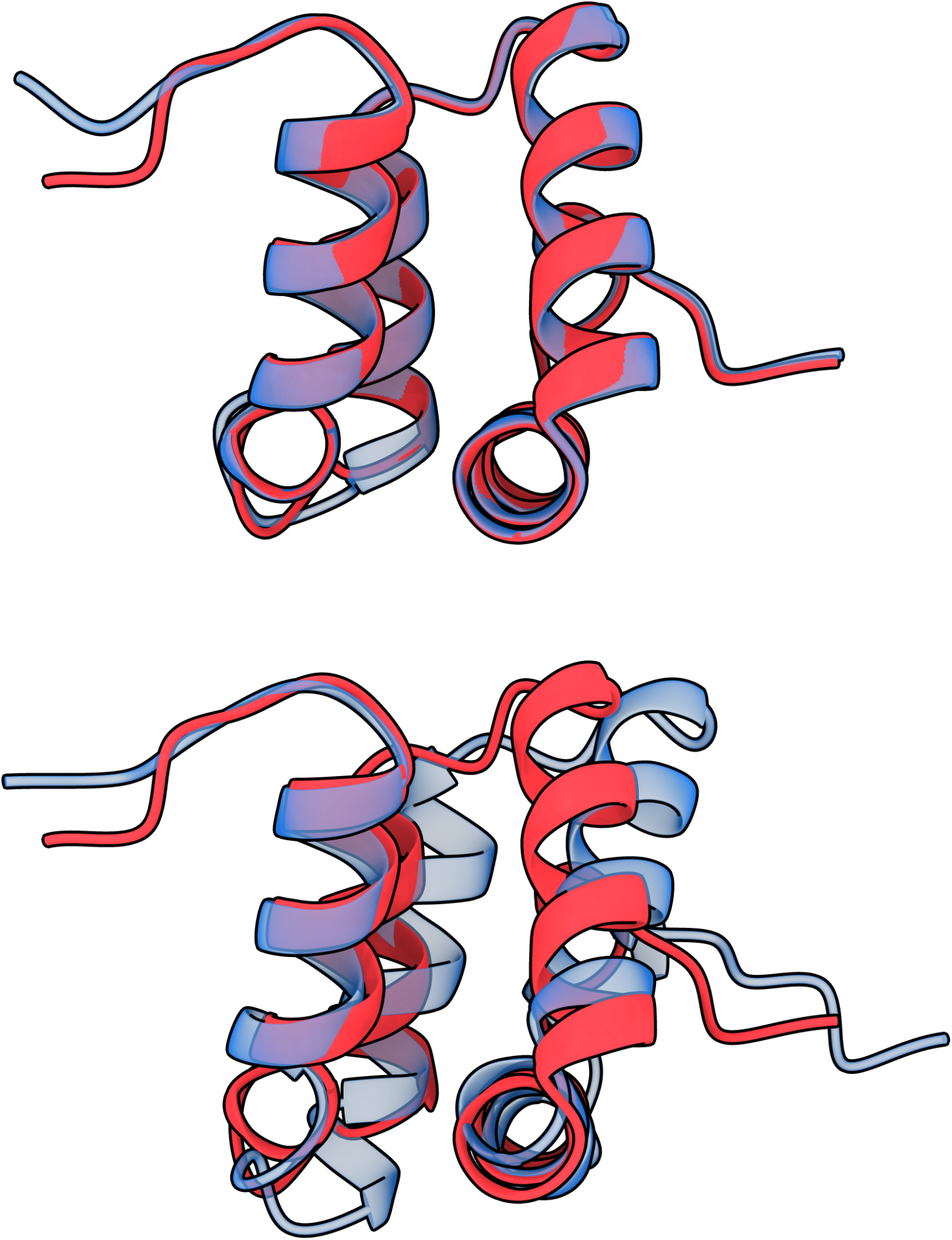
AlphaFold 2 predicts that mutations at the sites involved in the WLYY motif strongly influence tertiary structure without changing secondary structure more than mutations at random sites. AlphaFold 2 predicted tertiary structures for AVR3a 2A sequences: 1) wild type - W at site 36, Y at site 69 (solid red), 2) mutant with Y at site 36, W at site 69 (transparent blue, top), 3) mutant with K at sites 36 and 69 (transparent blue, bottom). Swapping the residues at sites 36 and 69 preserves the hydrophobic interactions that keep two helices together and results in a nearly unchanged tertiary structure (TM-align score [20] vs. the WT: 0.98). In contrast, replacing them with similarly charged hydrophilic residues that repel each other pushes the helices apart and results in a structure that diverges much more strongly from the WT (TM-align score: 0.64). Image created with UCSF ChimeraX version 1.10 [30].

### 3.6 The protein secondary structure map displays simplicity bias

Applying algorithmic information theory to genotype-to-phenotype maps has demonstrated that many of them, e.g., that of RNA secondary structure [31], are biased towards simple outputs [32]. More precisely, the a priori probability *P* (*x*) that randomly sampled inputs generate an output *x* decays exponentially with the approximate Kolmogorov complexity 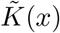 of that output:

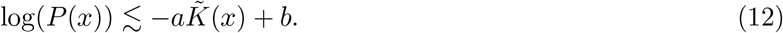

As far as we are aware, it has not been tested whether the *protein* secondary structure genotype-to-phenotype map exhibits simplicity bias. We characterize the complexity of secondary structure strings by calculating their Lempel-Ziv complexity, which has been shown to be an adequate approximation to the Kolmogorov complexity [32]. We use a definition analogous to that in [33]:

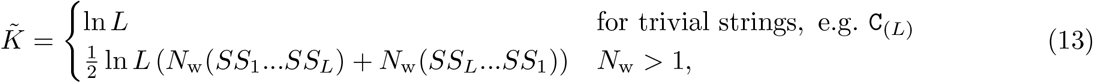

where *N*_w_(*SS*) is the number of words a string *SS* is parsed into according to the 1976 version of the Lempel-Ziv compression algorithm. We calculate *N*_w_ via a version of the algorithm by Kaspar and Schuster [34] and, following Dingle et al. [32], take the average of the secondary structure string *SS* and its reverse to obtain a finer-grained measure of complexity. We treat the case of *N*_w_ = 1 separately to distinguish between trivial strings of different lengths [32].

In Figure 4, we show that the upper bound of the logarithm of estimated genotype frequency scales linearly with the approximate Kolmogorov complexity 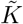, i.e., the protein secondary structure map displays simplicity bias. Note, however, that a more rigorous test for simplicity bias would require studying the secondary structures associated with randomly sampled genotypes.

**Figure 4.**
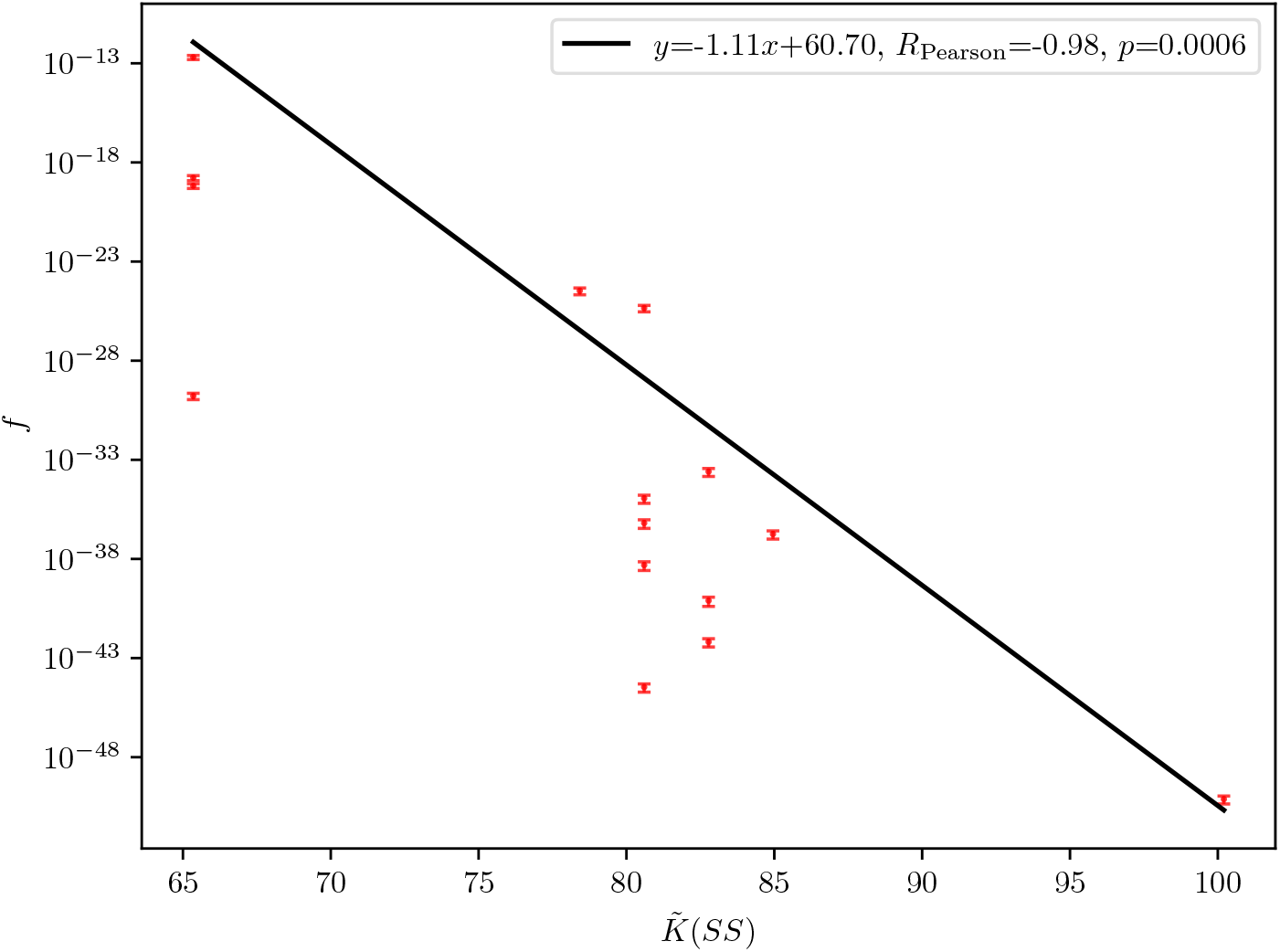
The protein secondary structure map exhibits simplicity bias: the logarithm of the upper bound for the frequency *f* of a secondary structure phenotype *SS*, fitted with a straight line (black) scales with the approximate Kolmogorov complexity of the phenotype 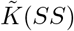. Thus, the phenotype estimated to be the most robust (⟨*r*⟩ = 0.689) is C_(4)_H_(72)_C_(2)_, where subscripted indices indicate the number of consecutive sites with the same secondary structure, has the longest continuous *α*-helix out of all studied here. We use phenotype frequency estimates calculated from exploring the point substitution mutants accessible via a point mutation in DNA for 10 sets of randomly chosen subsamples of 20 neutral genotypes. Error bars calculated based on the estimated standard error of the mean for the neutral component size, see Eq. (S2) in the SI.

The genotype estimated to be most robust, C_(4)_H_(72)_C_(2)_, has the longest unbroken helix out of those characterized here and a Kolmogorov complexity equal to that of the next two phenotypes with the highest ⟨*r*⟩, C_(5)_H_(70)_C_(3)_ and C_(4)_H_(71)_C_(3)_. This is in line with our observation from Section 3.2 that the sites most sensitive to mutation are those at or near the boundaries of structured regions: besides being simple, genotypes like these have fewer such sites than the WT.

### 3.7 Phenotype robustness scales logarithmically with phenotype frequency

Through studying mutated phenotypes we find that many of those that are very rarely encountered while exploring the neutral component of the 2A phenotype have a similar or higher robustness than it - see Table 1 and Figure 5. Although the frequencies of the phenotypes we have characterize within the dataset of 2A-derived mutants varies by ~ 3 orders of magnitude, their robustness ranges between 0.2 and 0.7, and is predicted to be highest for the phenotype C_(4)_H_(72)_C_(2)_, which is seen in only 0.15% of genotypes encountered in the scanning of the 2A neutral component. For more details on the relationship see Table 1 and Supplementary Figure S6. We find that the estimated robustness of these phenotypes varies logarithmically with their frequency over a range of more than 24 orders of magnitude for the latter. This logarithmic scaling has been observed in many unrelated genotype-to-phenotype maps and is thought to be one of their fundamental properties [19].

**Figure 5.**
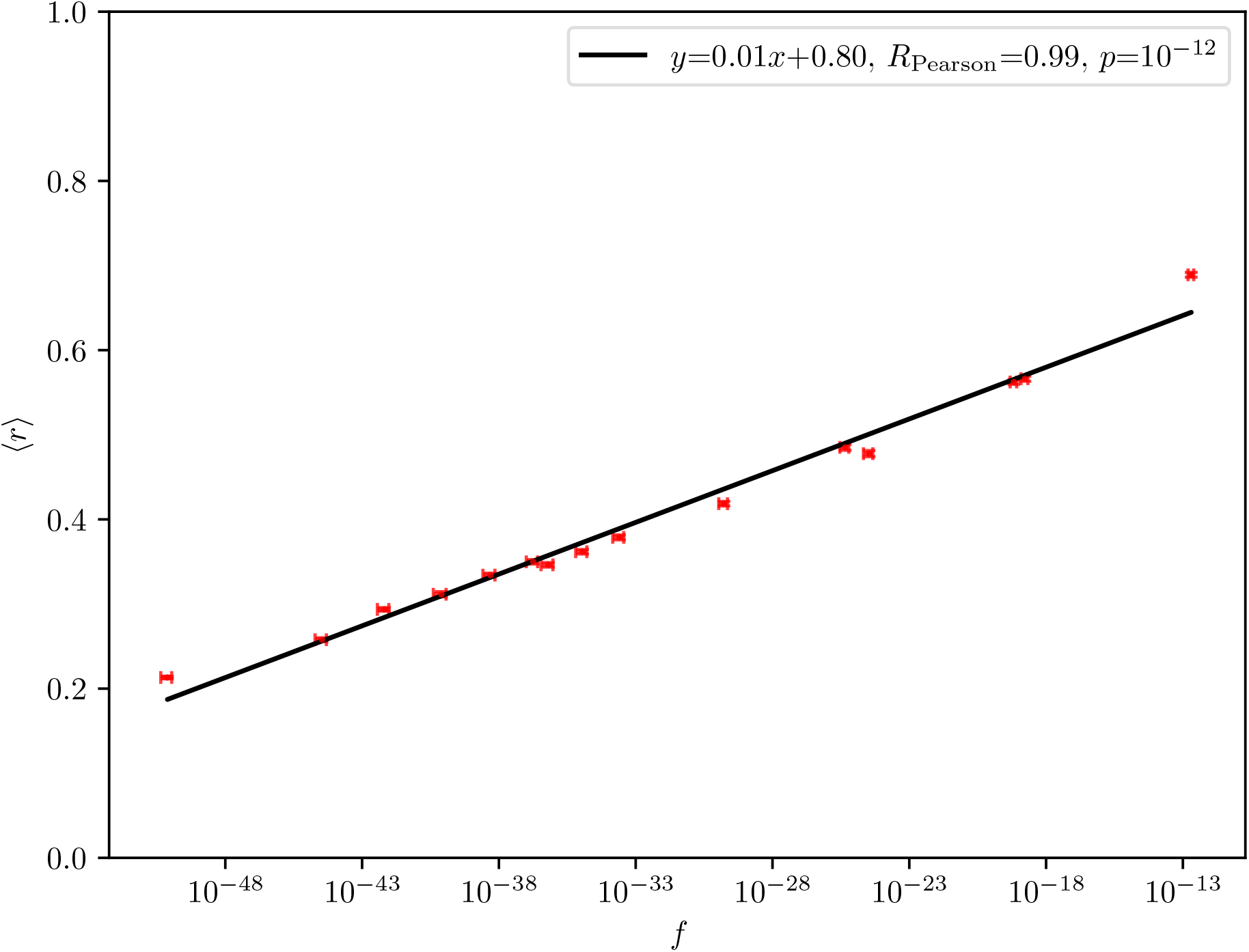
The phenotype robustness ⟨*r*⟩ scales logarithmically with frequency *f* across many orders of magnitude for *f*, as reported for many other GP maps [1, 19]. We use phenotype robustness and frequency estimates calculated from exploring the point substitution mutants accessible via a point mutation in DNA for 10 sets of randomly chosen subsamples of 20 neutral genotypes. Error bars calculated based on the estimated standard error of the mean for the neutral component size, see Eq. (S2) in the SI.

## 4 Conclusions and discussion

In this study, we have characterized the protein secondary structure genotype-to-phenotype map by using the machine-learning-based tool Porter 5 [6] to predict the secondary structure of mutants generated from the 2A variant of the *P. palmivora* AVR3a effector domain.

We have used site-scanning [2] to estimate the robustness of the secondary structure of this protein and its site-specific equivalent ⟨*F*^(k)^⟩. We have shown that a set of random subsamples of ~20 neutral genotypes from 10 independent runs allow calculating an adequate estimate of these properties for this protein of length 78 amino acids (Figure 1).

Our analysis indicates that sites at or close to the ends of structured regions tend to be most sensitive to mutations (Figure 2). We have also found that the residues found at these sites in our algorithmically generated mutants overwhelmingly tend to be those that preferentially form the secondary structure type predicted for that site in the WT protein. We find that sites 58-63 of the 2A AVR3a sequence are most sensitive to mutations with respect to secondary structure. In contrast with ‘phenotypic cliffs’ [35] seen in the RNA secondary structure GP map, we observe that mutations at the most sensitive sites tend to be the ones that cause the greatest phenotypic changes.

Through a small-scale AlphaFold 2 study we show that mutations at sites 36 and 59, both identified as sensitive via our Porter 5 study, change both the tertiary and the secondary structure of the protein significantly more than point mutations at random other sites. In contrast, mutations at site 1, which our Porter 5 study identifies as robust, change neither the secondary nor the tertiary structure of the protein significantly more than point mutations at a random other site. AlphaFold 2 predictions, however, predict that site 61 is similarly robust, in contrast with our results obtained via Porter 5.

Our analysis of site sensitivity with respect to secondary structure identifies only two of the four sites that form the conserved WLYY motif in AVR3a effector proteins as relatively sensitive to mutation. This is not surprising considering that hydrophobic interactions between these sites maintain a specific spatial arrangement of the helices in the protein [25], meaning that they contribute to its tertiary structure. We show that AlphaFold 2 predicts that mutations at these sites, particularly the introduction of similarly charged amino acids, disproportionately affect tertiary structure without generally changing secondary structure more than mutations at random other sites (Figure 3).

We have characterized a set of mutated phenotypes encountered during the exploration of the 2A AVR3a secondary structure phenotype. These range from secondary structures very rarely found in the 2A run to some of the most common ones (see Table 1, Figure S6). We estimate that the robustness of all of these is similar to that of the WT phenotype despite the large difference in the number of genotypes that map to them within the set generated through mutating the 2A sequence.

Moreover, we observe that phenotypes with long uninterrupted *α*-helices than are more stable than those with shorter helical regions separated by coil residues. Since such phenotypes are algorithmically simpler, this observation is in line with the previously reported simplicity bias in other genotype-to-phenotype maps such as that of RNA secondary structure [31, 32], as seen in Figure 4.

Finally, we observe that the robustness of the studied phenotypes is proportional to the logarithm of their frequency across many orders of magnitude for the latter (Fig. 5), in line with what has been reported for other genotype-to-phenotype maps [1, 19].

A caveat to our results is that, since we do not have an accurate estimate for the likelihood of a given phenotype, we consider only mutations that map to the same phenotype as the WT to be neutral. It is possible to conduct a more nuanced analysis of GP maps that can be approximated by algorithms that calculate the probabilities of different phenotypes, such as that of RNA secondary structure [35].

Another limitation of our study is that it uses a machine-learning based secondary structure prediction algorithm that uses alignment to known proteins. It is unclear how accurate such algorithms are for sequences that are not similar to ones used for their training, and even the effect of point mutations is known to be difficult to predict with such tools, e.g., AlphaFold 2 [36]. At the same time, one may expect that algorithms trained on existing protein structure databases such as Porter 5 give predictions that are sufficiently accurate to describe the basic properties of real-world protein genotype-to-phenotype maps.

Similar algorithmic approaches can be used to design mutated proteins with preserved secondary and/or tertiary structure. Such proteins may have advantages such as a more desirable immune response through altered interactions with antibodies. Furthermore, neutral mutants could be algorithmically designed to be more efficiently produced in vivo through accounting for codon bias. Further exploration of the map of genotype-to-phenotype maps such as that of protein secondary structure holds promise for yielding insights into the evolution of a wide range of biological systems, important examples being pathogens such as the oomycete *P. palmivora* and cancer cells.

## 5 Acknowledgments

This work was supported by the Isaac Newton Trust (NQAG/341) and the University of Cambridge School of Technology Seed Fund and was performed using resources provided by the Cambridge Service for Data Driven Discovery (CSD3) operated by the University of Cambridge Research Computing Service (www.csd3.cam.ac.uk), provided by Dell EMC and Intel using Tier-2 funding from the Engineering and Physical Sciences Research Council (capital grant EP/T022159/1), and DiRAC funding from the Science and Technology Facilities Council (www.dirac.ac.uk). JKN acknowledges funding from the MRC (Grant MC FE 00035) Cross Disciplinary Fellowship (XDF) Programme.

