## Supplementary Material for "Exploring the large-scale properties of a protein secondary structure genotype-to-phenotype map"

#### 1 Additional methods

We measure the average number of changes to the secondary structure induced by a point mutation for each site, $\langle N^{(k)}_{\mathrm{SS\ changes}}/N^{(k)}_{\mathrm{mut\ 0}}\rangle$, where $N^{(k)}_{\mathrm{mut\ 0}}$ is the number of point amino acid mutations accessible via a point mutation in DNA; $N^{\mathrm{k}}_{\mathrm{mut\ 0}}$ depends on the genotype that is being mutated.

We calculate finer-grained measures of the secondary structure changes induced by mutations from the scores assigned by Porter 5 to each type of secondary structure at each site. These scores, $p^{(k)}_{\mathrm{helix}}$, $p^{(k)}_{\mathrm{sheet}}$ and $p^{(k)}_{\mathrm{coil}}$, add up to unity at each site 'k' for a given predicted secondary structure. We calculate a function that measures the average effect of a mutation at site 'k' on site 'l' as follows:

$$\langle P^{(\mathrm{kl})}_{\mathrm{diff}}\rangle = \frac{1}{2}\sum_{\mathrm{i=1}}^{N_{\mathrm{samples}}}\sum_{\mathrm{j=1}}^{N_{\mathrm{subsample}}}\frac{1}{N^{(k)}_{\mathrm{ij\ mut}}}\left(\left|p^{(l)}_{\mathrm{helix\ mut}}-p^{(l)}_{\mathrm{helix\ WT}}\right|+\left|p^{(l)}_{\mathrm{sheet\ mut}}-p^{(l)}_{\mathrm{sheet\ WT}}\right|\right), \tag{S1}$$

where $P^{(\mathrm{kl})}_{\mathrm{diff}} = 1$ would mean that mutating site 'k' introduces the maximum possible change in the predicted score for site 'l' in all studied mutants, whereas at $P^{(\mathrm{kl})}_{\mathrm{diff}} = 0$ the mutation induces no change in the predicted scores. We show a plot of this function for the 2A phenotype in Figure S1 and all other phenotypes in Figure S5.

Comparison of Figure 2 and Figure S1 indicates that mutations at some sites in genotypes with the AVR3a phenotype, particularly 35-36 and 59, affect the secondary structure at a large number of sites, including distant ones. This predicted non-local effect is especially strong at site 59, which as we discuss in the main text allows very little sequence variation. Point mutations there on average lead to the highest number of secondary structure changes and are predicted to have a strong influence the secondary structure at sites 30-35.

We use a common approximation for error propagation to estimate the pooled error of the mean of the

neutral component size $S_{\mathrm{NC}}$ [1], for which we use the estimate in Eq. (10):

$$\hat{\sigma}_{\mathrm{pooled}\ S_{\mathrm{NC}}} \approx \sqrt{\sum_{\mathrm{k}=1}^{\mathrm{k}=L} \left( \hat{\sigma}_{\mathrm{pooled} \langle F^{(\mathrm{k})}_{\mathrm{neutr\ subsample}} \rangle} \partial_{\langle F^{(\mathrm{k})}_{\mathrm{neutr\ subsample}} \rangle} S_{\mathrm{NC}} \right)^2} = S_{\mathrm{NC}} N_{\mathrm{mut}} \sum_{\mathrm{k}=1}^{\mathrm{k}=L} \sqrt{\left( \frac{\hat{\sigma}_{\mathrm{pooled} \langle F^{(\mathrm{k})}_{\mathrm{neutr\ subsample}} \rangle}}{(1 + N_{\mathrm{mut}} \langle F^{(\mathrm{k})}_{\mathrm{neutr\ subsample}} \rangle)} \right)^2}. \tag{S2}$$

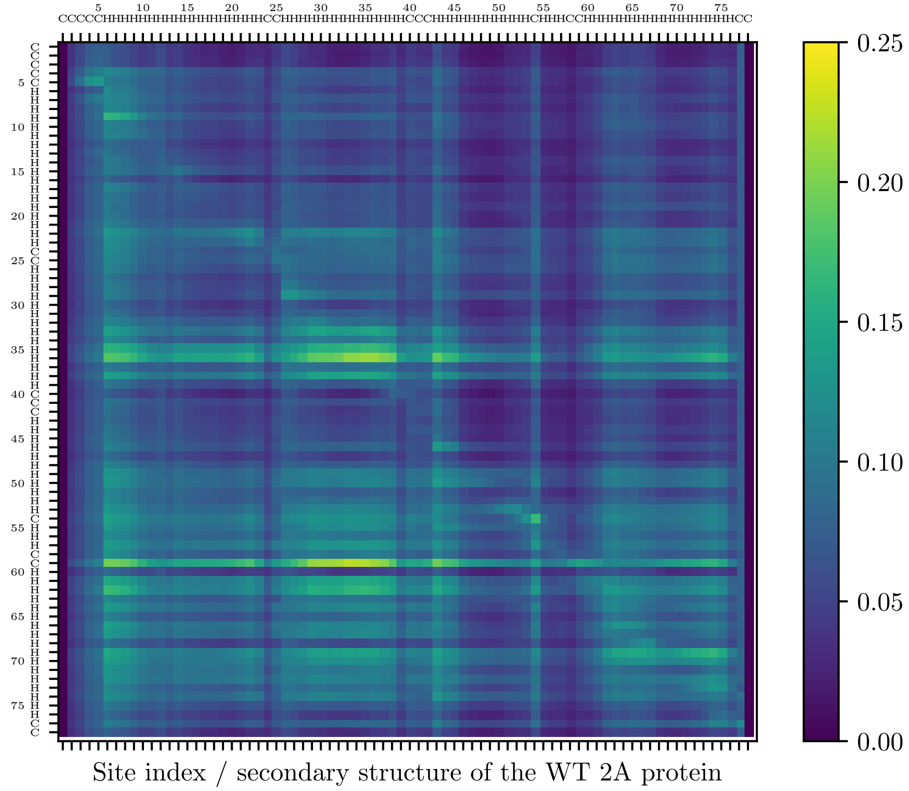


Figure S1: **Matrix measuring the average effect of mutating site 'k' on site 'l' on the 2A AVR3a effector domain secondary structure phenotype, $\langle P^{(\mathrm{kl})}_{\mathrm{diff}} \rangle$ in terms of the scores calculated by Porter 5 that measure its estimate for the probability of helix or sheet.** Mutations at several sites, particularly 36 and 59, have a substantial effect on the secondary structure at other sites, seen as horizontal streaks with high $\langle P^{(\mathrm{kl})}_{\mathrm{diff}} \rangle$ values.

### 2 Supplementary data and code

We provide example scripts for data generation and analysis, as well as sequence data for the studied phenotypes and the top-rated AlphaFold 2 predictions for the proteins whose tertiary structure we discuss in the paper.

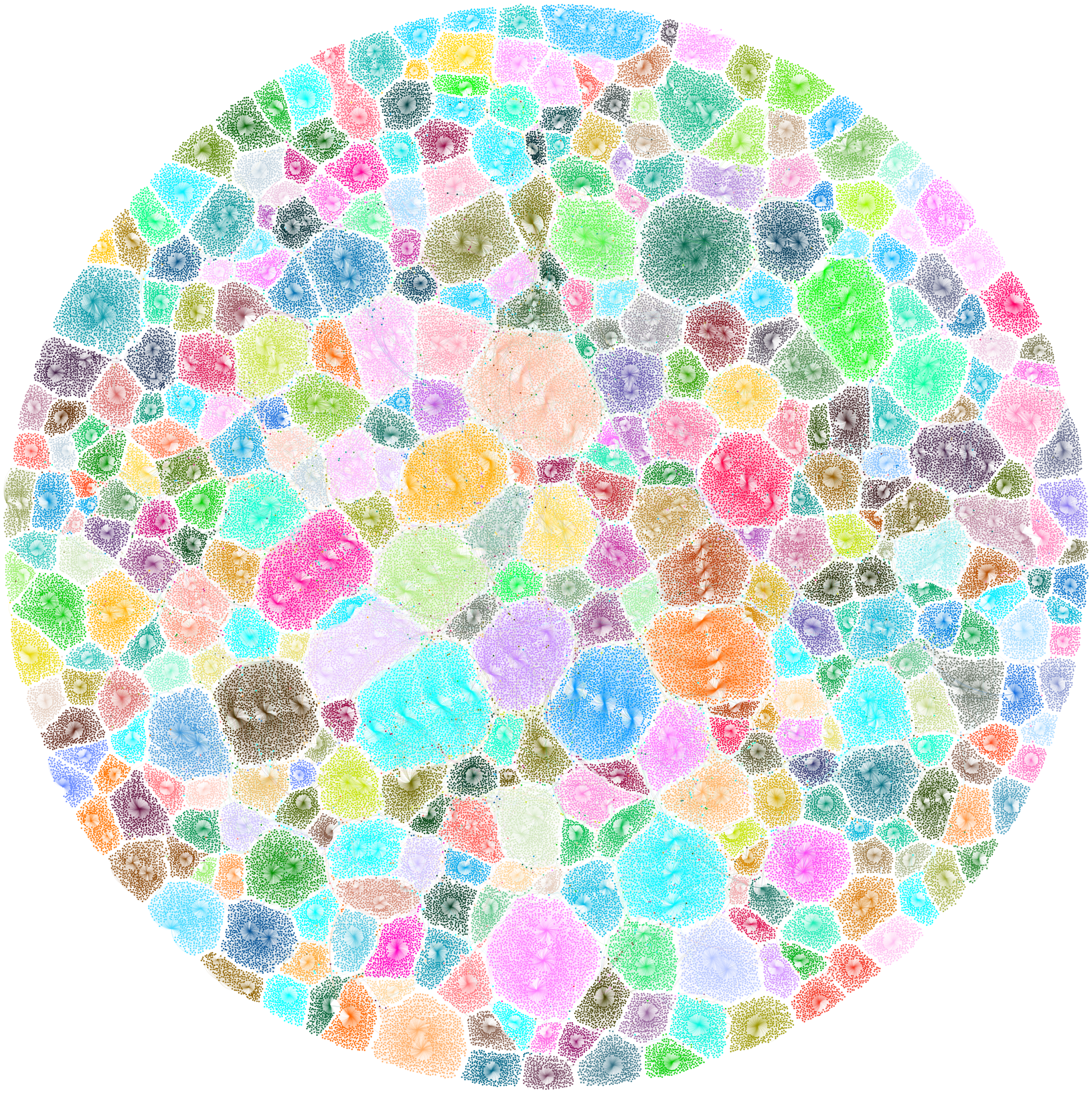

Figure S2: **Qualitative visualization of the part of the neutral component of the 2A AVR3a effector domain SS phenotype explored in this study.** Network diagram of all genotypes generated via site-scanning (see Methods) predicted to have the same secondary structure phenotype as the 2A AVR3a sequence. Colours correspond to communities determined via the Louvain algorithm as implemented in Gephi [2].

| 'WT' sequence | Frequency rank (2A dataset) | Sequence Hamming distance from the WT |
|---|---|---|
| TPSFNLAQLTKSKHAAALAEQLMGDHKLADAAYIWWQHNGVTLTKLDDFLKLASRKTQGRKYDQIYNAYMMHLGYTGV | 1 (WT) | 0 |
| MKFFRVSKLRISEQVNDWTEQVFSGQTMVVGAYRWFIPKGVRMSQLYDVMKLASLKKQGKKSNQSFNDFVMLLEYMLA | 4 | $6.79 \times 10^{-1}$ |
| TPNFNLAQLTKSRNAAALDEQLMGDHKLSDAAYIWWQHNGVTLTKLDDFLKLASRKAQGRKYDQIYNAYMMHMGYTGV | 6 | $8.97 \times 10^{-2}$ |
| SASIGMGFVNKLNMEARALVHLFCDHKSVTYSFSWWEHKFVENAKVRIFTVYASRKARGQRFNTYHFSYMLHIGYIGV | 9 | $6.79 \times 10^{-1}$ |
| TMRFNTAWITKEKDKSKVAAKLMGNHKVVGCACVCGRKYDLECKHKVDFMRIASIKLRGQQYTQRTNACIFKIVNISV | 12 | $6.79 \times 10^{-1}$ |
| GAVLHWTEVANSWNLSRIGATMRIDEVLWFERYNWWQHVCATLIVDFDFLKRVNQRTKGWKFDCIYNAVMMHIGFVRR | 15 | $6.67 \times 10^{-1}$ |
| WSMLNLEYVNQIELAQAAKKQINSNDHCIDMIFMIWRRNCIPITEAVTHLILASVLTIGPKYKQICQTYILVLGLTGK | 17 | $6.92 \times 10^{-1}$ |
| TRAFDEERTSLRDLQIALVHKVTMTIDQSCAAYMWIEFNGVYLKSIHGILRSASRRTEGLKNQQLYSALAFCLARTGG | 24 | $6.79 \times 10^{-1}$ |
| VPRLPLAFLIKMKWTGQMTSLVMDMKPVARQAHRWWYQKYSEVRSFDNYFRLVNRKSIGESYFQFYNNYMQQIKCACF | 34 | $7.18 \times 10^{-1}$ |
| SILFHVTTKAIIADIHSSLEQLYAKDTLRAFVNQWCFILCLRVNKYHDILKIESNQMQGSKYLQIYDEYMMSLAFTFD | 54 | $7.05 \times 10^{-1}$ |
| QFIFHRIDSTKDEWAVRMFERLNANLPSIYAIYIWYDHNGTSCIRLNTIFNEVHRSTQGKKFDVMYQAFMLYLGDFYY | 75 | $6.79 \times 10^{-1}$ |
| TRRYKMAGKTEAQDKLKTATEFMGDDKLARCERIFVMMYWLDFQLQGEFLWLASQKIQGPHYNKCSNAYVFNIANLSE | 101 | $7.05 \times 10^{-1}$ |
| SILFHVTTKAIIADIHSSLERLYAKDTLRAFFYQLYFILCLRVNKYHDILKIVSNQMRGSTYLQIYNEYMMSLAFTFD | 105 | $7.31 \times 10^{-1}$ |
| TRTYKMAGKTEAQAKLKTATEFMGDDKLASCERIYVMMYWLDFQLQGEFLWLASQRRQGPHYNKCSNAYVFIIANLSV | 117 | $7.05 \times 10^{-1}$ |
| TMCFNTAWITKEIDKSTVAAKLMGTHKVAGCACVCGRKFDLEYKHKVDFMRIASIKMRGQQYTQRTNACIIKIVNISV | 244 | $6.79 \times 10^{-1}$ |

Table S1: Additional data on the mutated phenotypes from Table 1 selected for further study: starting sequences for site-scanning and its Hamming distance from the 2A AVR3a effector domain sequence.

Figure S3: *(following and current pages)* **Fraction of neutral mutations, $\langle F^{(\mathrm{k})}\rangle$ and normalized number of secondary structure changes per non-neutral mutant, $\langle \tilde{N}^{(\mathrm{k})}_{\mathrm{SS\ changes}}\rangle$, for each site in various phenotypes.**

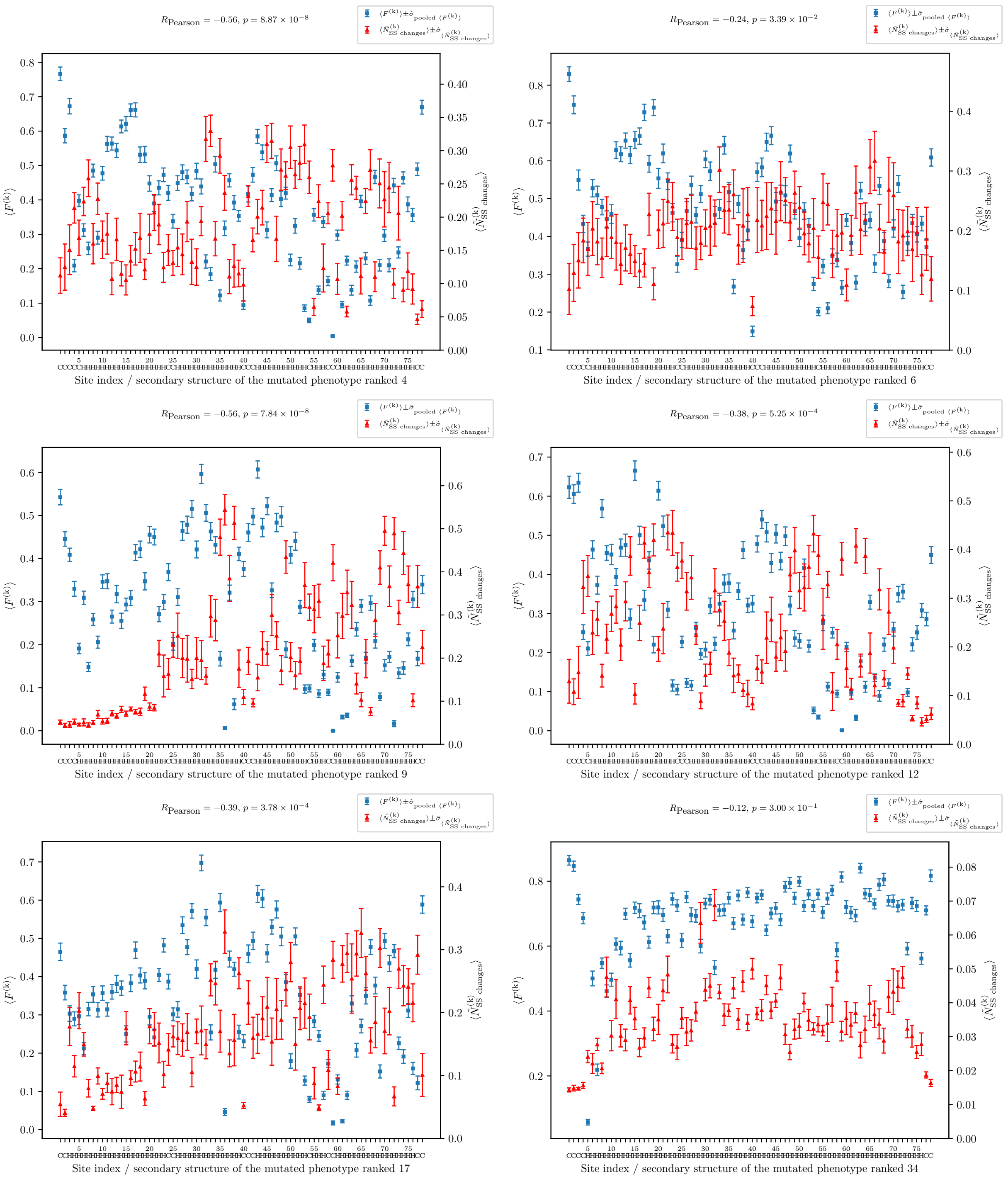

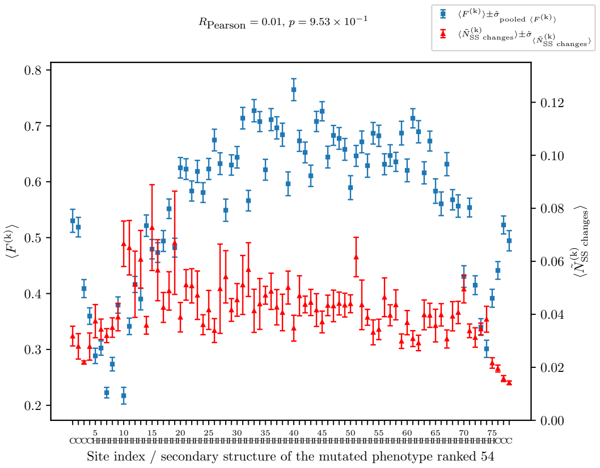

$R_{\text{Pearson}} = 0.01$, $p = 9.53 \times 10^{-1}$
$\langle F^{(k)} \rangle \pm \hat{\sigma}_{\text{pooled } \langle F^{(k)} \rangle}$
$\langle \tilde{N}^{(k)}_{\text{SS changes}} \rangle \pm \hat{\sigma}_{\langle \tilde{N}^{(k)}_{\text{SS changes}} \rangle}$
$\langle F^{(k)} \rangle$
$\langle \tilde{N}^{(k)}_{\text{SS changes}} \rangle$
Site index / secondary structure of the mutated phenotype ranked 54

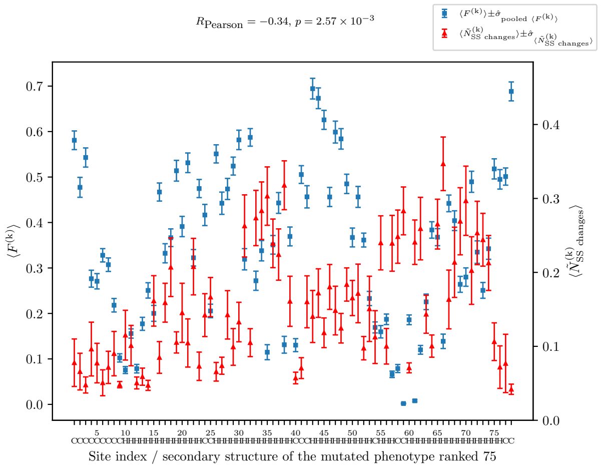

$R_{\text{Pearson}} = -0.34$, $p = 2.57 \times 10^{-3}$
$\langle F^{(k)} \rangle \pm \hat{\sigma}_{\text{pooled } \langle F^{(k)} \rangle}$
$\langle \tilde{N}^{(k)}_{\text{SS changes}} \rangle \pm \hat{\sigma}_{\langle \tilde{N}^{(k)}_{\text{SS changes}} \rangle}$
$\langle F^{(k)} \rangle$
$\langle \tilde{N}^{(k)}_{\text{SS changes}} \rangle$
Site index / secondary structure of the mutated phenotype ranked 75

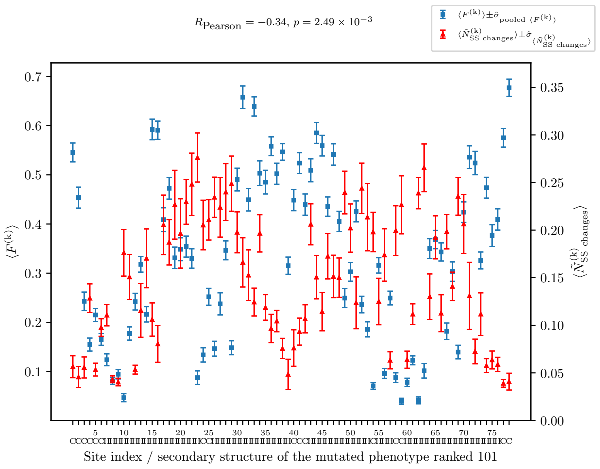

$R_{\text{Pearson}} = -0.34$, $p = 2.49 \times 10^{-3}$
$\langle F^{(k)} \rangle \pm \hat{\sigma}_{\text{pooled } \langle F^{(k)} \rangle}$
$\langle \tilde{N}^{(k)}_{\text{SS changes}} \rangle \pm \hat{\sigma}_{\langle \tilde{N}^{(k)}_{\text{SS changes}} \rangle}$
$\langle F^{(k)} \rangle$
$\langle \tilde{N}^{(k)}_{\text{SS changes}} \rangle$
Site index / secondary structure of the mutated phenotype ranked 101

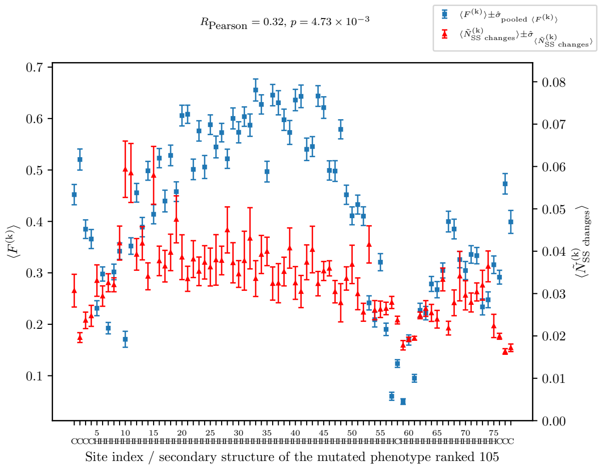

$R_{\text{Pearson}} = 0.32$, $p = 4.73 \times 10^{-3}$
$\langle F^{(k)} \rangle \pm \hat{\sigma}_{\text{pooled } \langle F^{(k)} \rangle}$
$\langle \tilde{N}^{(k)}_{\text{SS changes}} \rangle \pm \hat{\sigma}_{\langle \tilde{N}^{(k)}_{\text{SS changes}} \rangle}$
$\langle F^{(k)} \rangle$
$\langle \tilde{N}^{(k)}_{\text{SS changes}} \rangle$
Site index / secondary structure of the mutated phenotype ranked 105

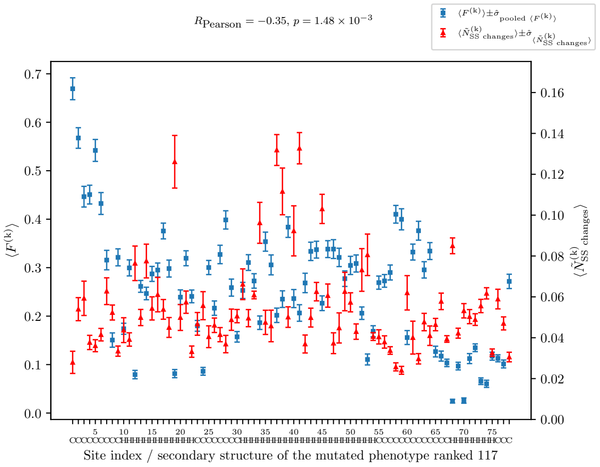

$R_{\text{Pearson}} = -0.35$, $p = 1.48 \times 10^{-3}$
$\langle F^{(k)} \rangle \pm \hat{\sigma}_{\text{pooled } \langle F^{(k)} \rangle}$
$\langle \tilde{N}^{(k)}_{\text{SS changes}} \rangle \pm \hat{\sigma}_{\langle \tilde{N}^{(k)}_{\text{SS changes}} \rangle}$
$\langle F^{(k)} \rangle$
$\langle \tilde{N}^{(k)}_{\text{SS changes}} \rangle$
Site index / secondary structure of the mutated phenotype ranked 117

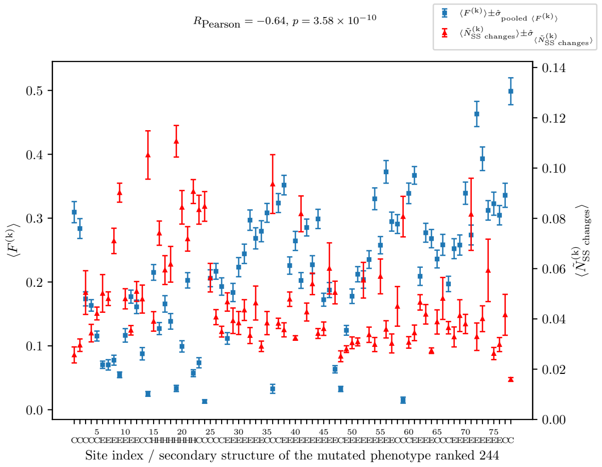

$R_{\text{Pearson}} = -0.64$, $p = 3.58 \times 10^{-10}$
$\langle F^{(k)} \rangle \pm \hat{\sigma}_{\text{pooled } \langle F^{(k)} \rangle}$
$\langle \tilde{N}^{(k)}_{\text{SS changes}} \rangle \pm \hat{\sigma}_{\langle \tilde{N}^{(k)}_{\text{SS changes}} \rangle}$
$\langle F^{(k)} \rangle$
$\langle \tilde{N}^{(k)}_{\text{SS changes}} \rangle$
Site index / secondary structure of the mutated phenotype ranked 244

Figure S4: *(following and current pages)* **The estimates for the robustness of the studied mutated phenotypes approximately converge at sparse subsample size $\sim 20$. Using a random subsample yields robustness intermediate between that estimated by choosing a subsample of neutral genotypes that are approximately equidistant from the WT sequence and that from subsamples of genotypes that are approximately equidistant in the number of site-scanning steps required to derive them from the WT.** The dashed line shows the estimate based on the fraction of genotypes encountered in the site-scanning run that map onto the WT phenotype.

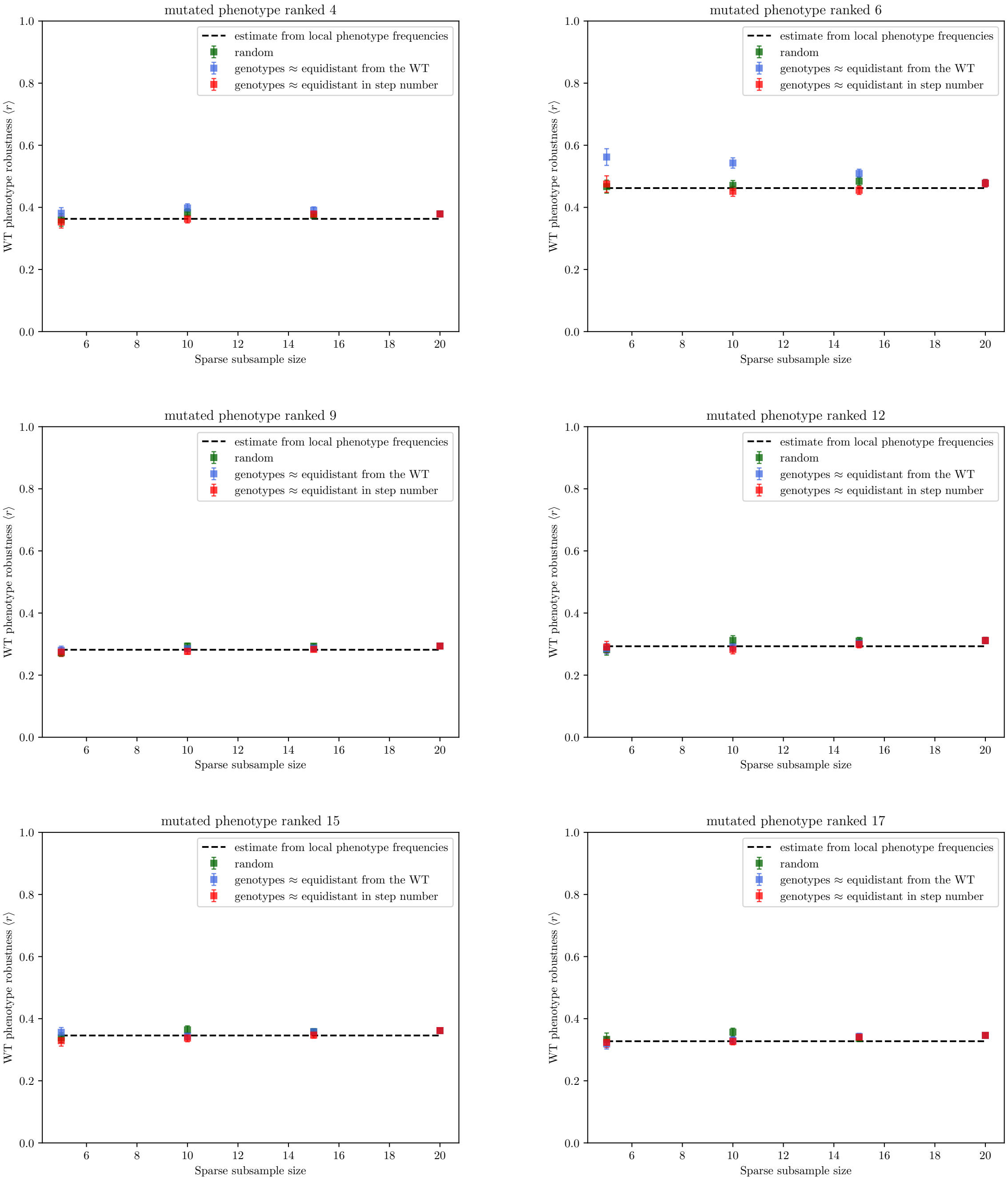
mutated phenotype ranked 4
mutated phenotype ranked 6
mutated phenotype ranked 9
mutated phenotype ranked 12
mutated phenotype ranked 15
mutated phenotype ranked 17
estimate from local phenotype frequencies
random
genotypes ≈ equidistant from the WT
genotypes ≈ equidistant in step number
WT phenotype robustness ⟨r⟩
Sparse subsample size
0.0
0.2
0.4
0.6
0.8
1.0
6
8
10
12
14
16
18
20

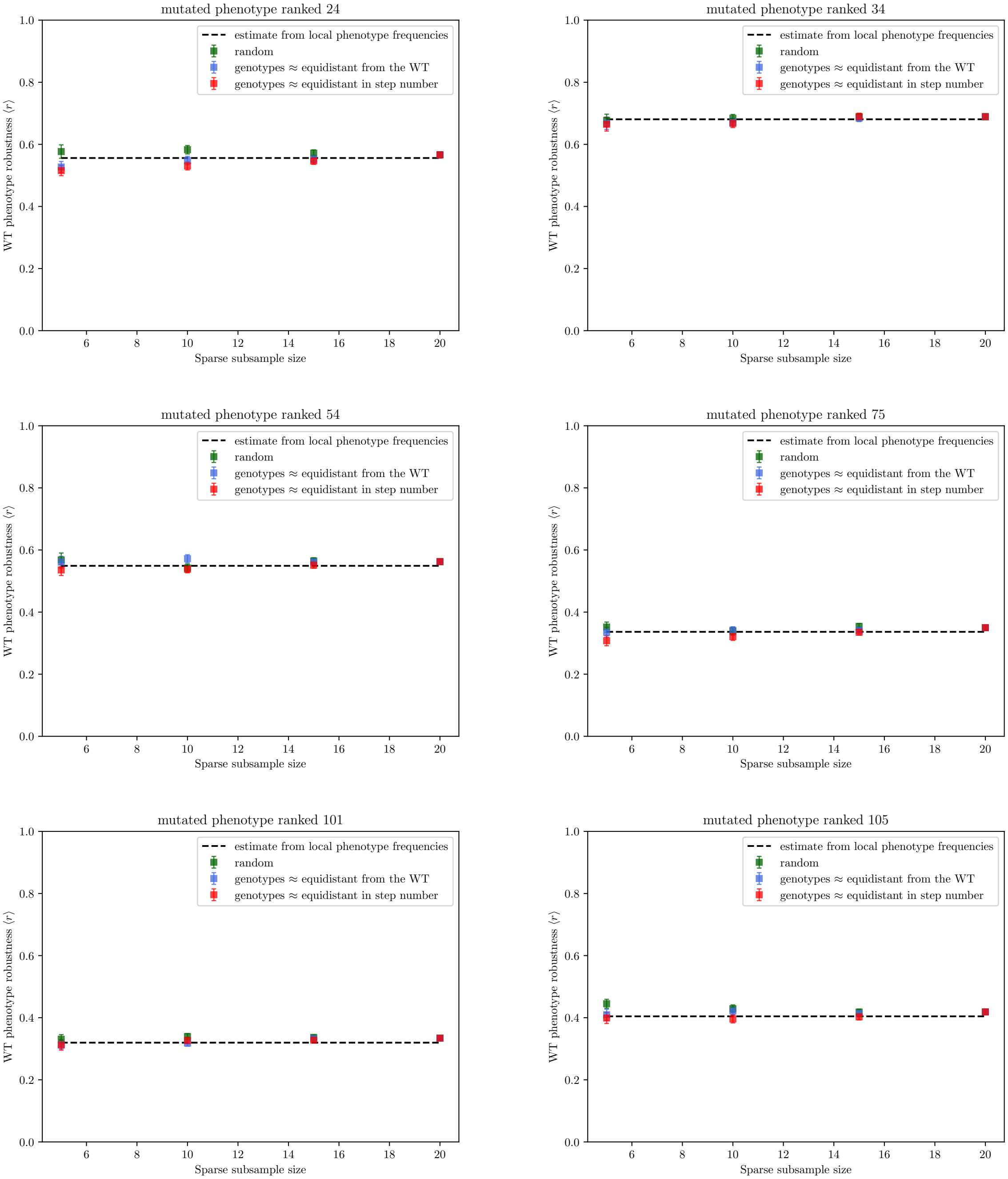
mutated phenotype ranked 24
mutated phenotype ranked 34
mutated phenotype ranked 54
mutated phenotype ranked 75
mutated phenotype ranked 101
mutated phenotype ranked 105
estimate from local phenotype frequencies
random
genotypes ≈ equidistant from the WT
genotypes ≈ equidistant in step number
WT phenotype robustness ⟨r⟩
Sparse subsample size

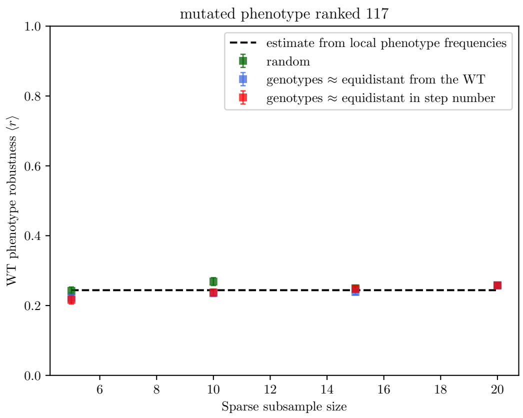


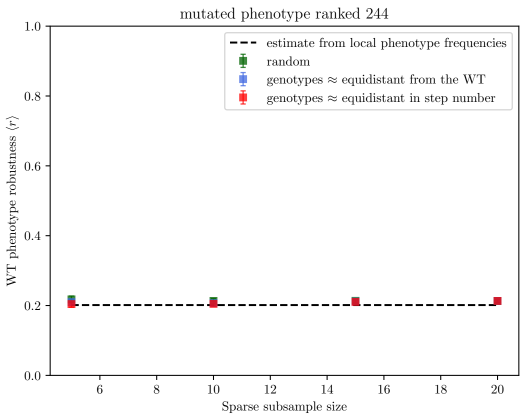


Figure S5: *(following pages)* **Matrices measuring the average effect of mutating site 'k' on site 'l' all studied mutated secondary structure phenotype, $\langle P_{\mathrm{diff}}^{(\mathrm{kl})} \rangle$. Mutations at some rows are predicted to have a strong effect on the secondary structure of other sites.**

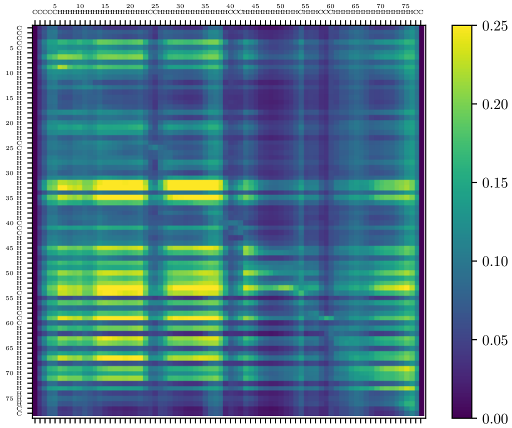

Site index / secondary structure of the mutated phenotype ranked 4

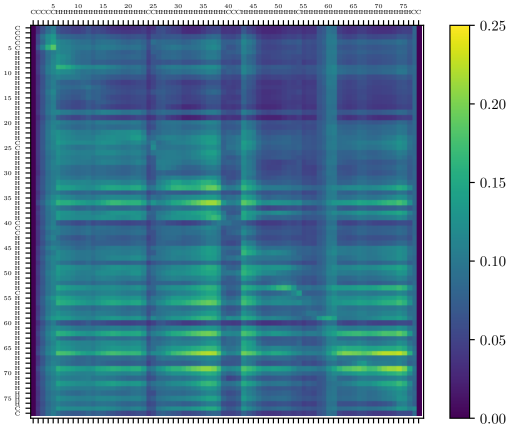

Site index / secondary structure of the mutated phenotype ranked 6

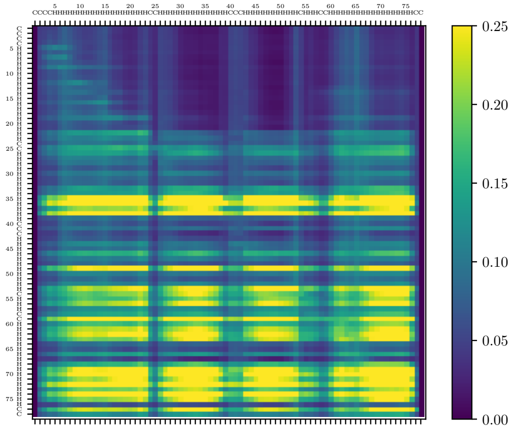

Site index / secondary structure of the mutated phenotype ranked 9

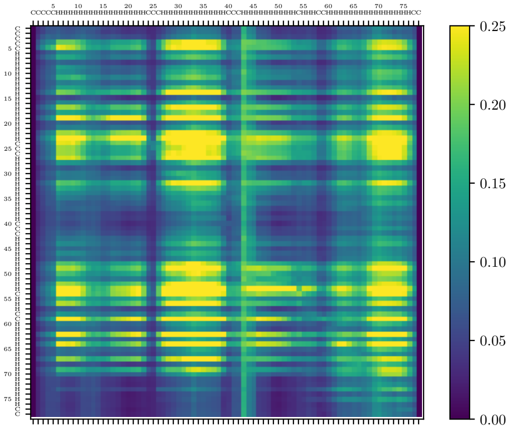

Site index / secondary structure of the mutated phenotype ranked 12

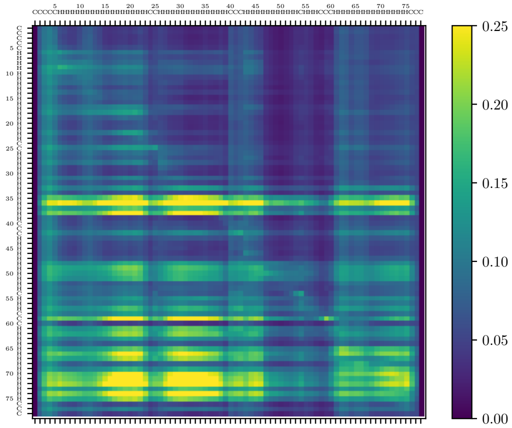

Site index / secondary structure of the mutated phenotype ranked 15

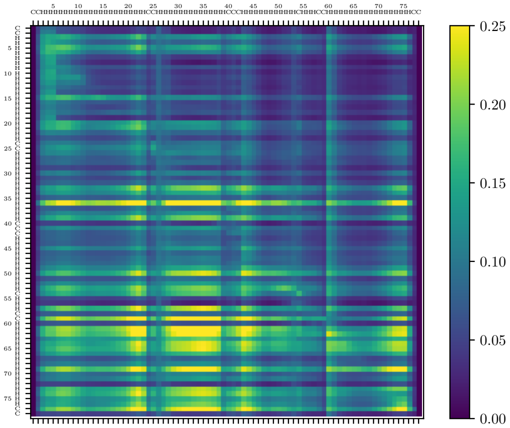

Site index / secondary structure of the mutated phenotype ranked 17

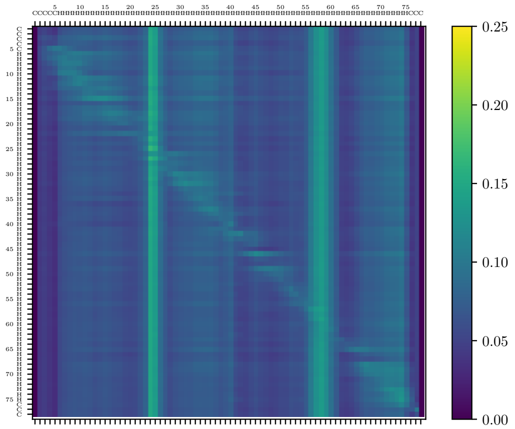

Site index / secondary structure of the mutated phenotype ranked 24

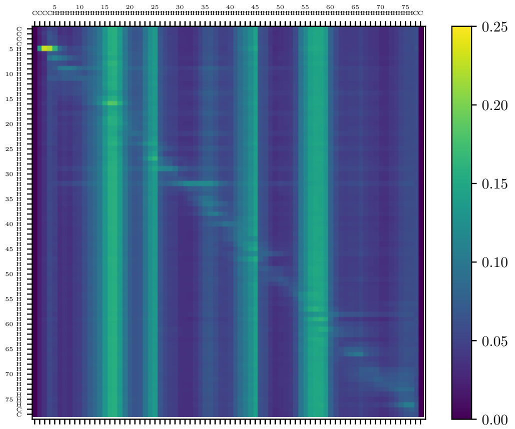

Site index / secondary structure of the mutated phenotype ranked 34

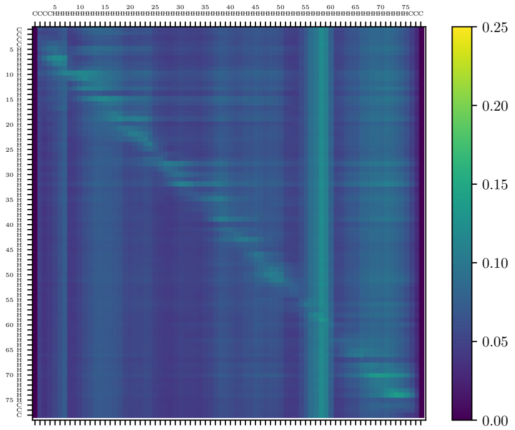

Site index / secondary structure of the mutated phenotype ranked 54

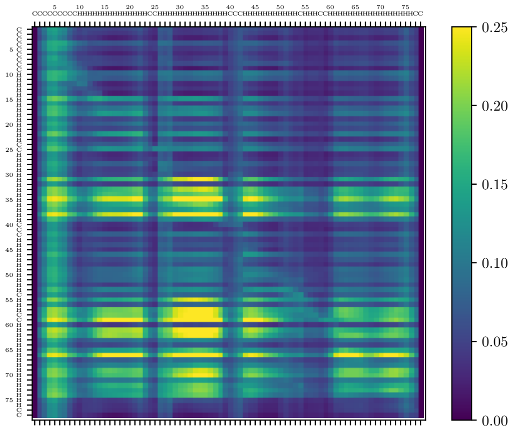

Site index / secondary structure of the mutated phenotype ranked 75

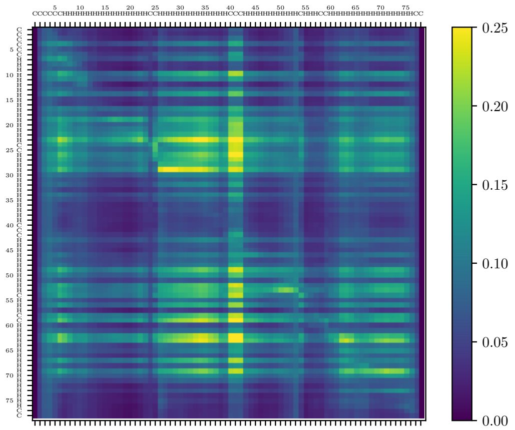

Site index / secondary structure of the mutated phenotype ranked 101

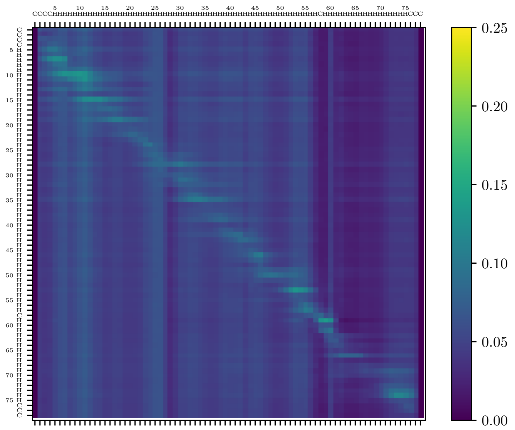

Site index / secondary structure of the mutated phenotype ranked 105

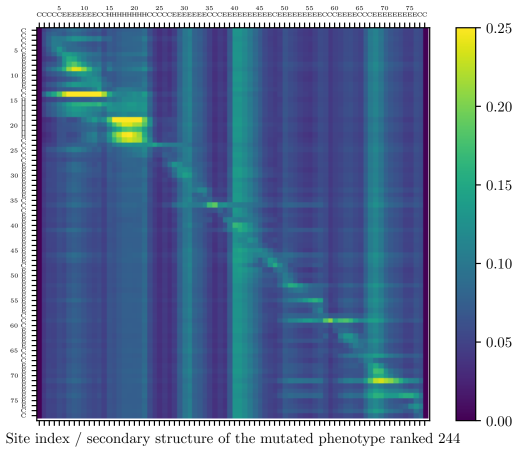

0.25
0.20
0.15
0.10
0.05
0.00
Site index / secondary structure of the mutated phenotype ranked 244

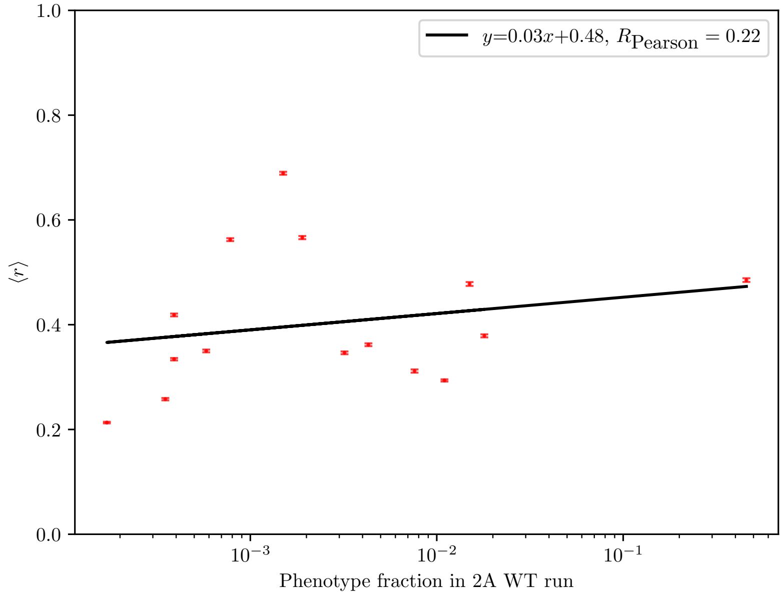


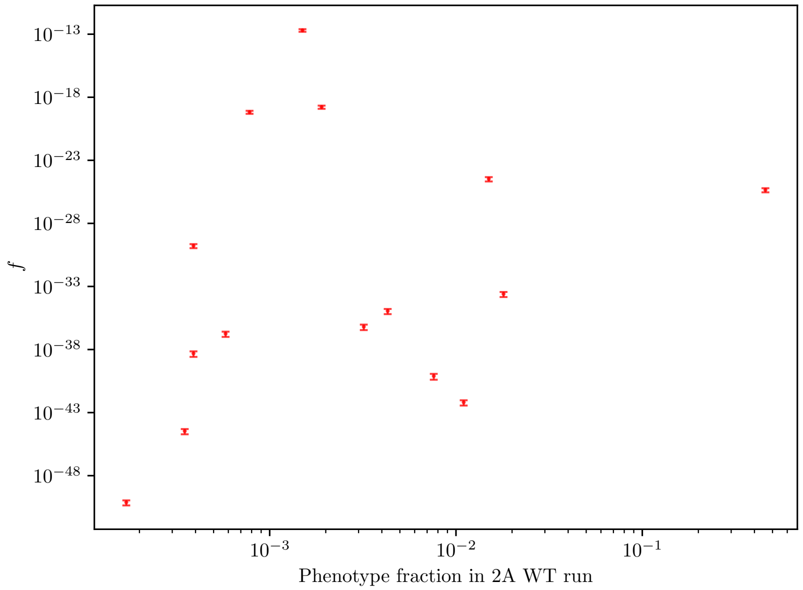


Figure S6: **Phenotypes that are rarely encountered while scanning the neutral component of the 2A phenotype are similarly robust to it.** The correlation between the fraction of genotypes mapping to these phenotypes within the 2A run (i.e., their local frequency in the 2A run) and their robustness (top) is negligible. The same is seen when plotting the estimated global phenotype frequencies of the same phenotypes versus their local frequencies (bottom). Error bars for the bottom plot calculated based on the estimated standard error of the mean for the neutral component size, see Eq. (S2) in the SI.

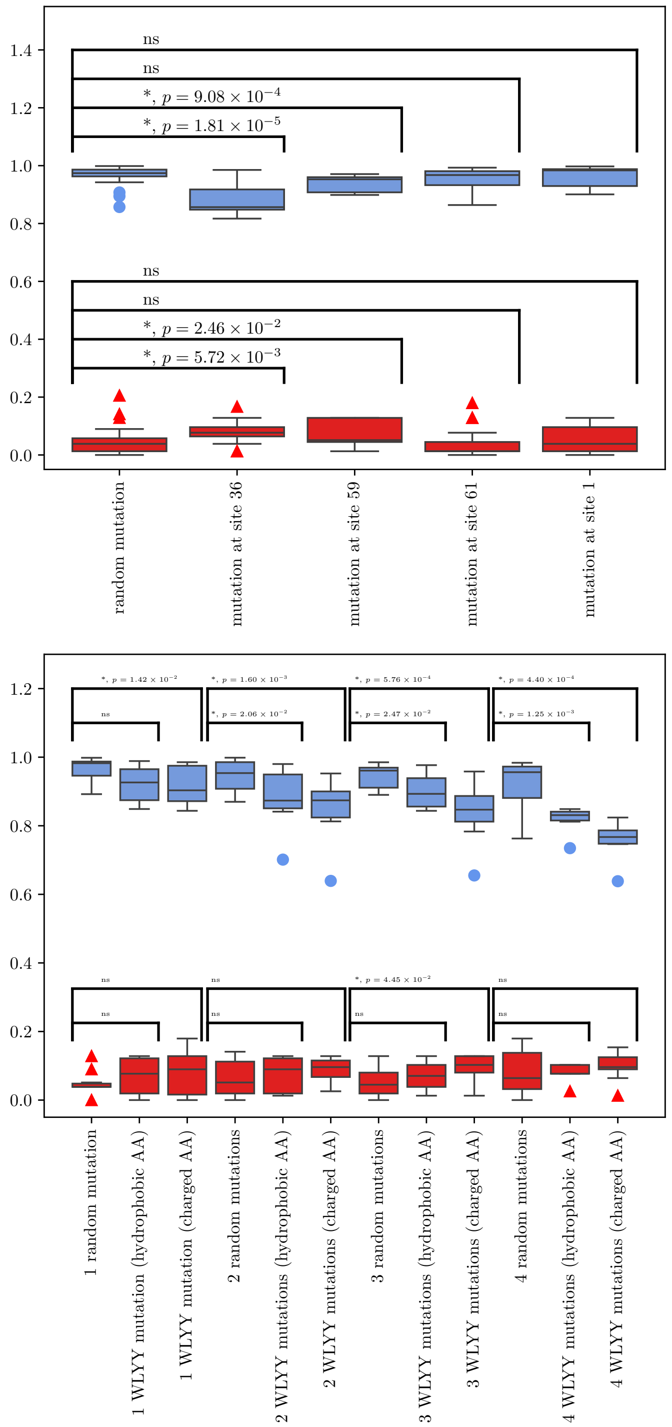


Figure S7: **Box-and-whisker plots showing TM-align scores (blue, outliers shown as circles) and normalized Hamming distances (red, outliers shown as triangles) from the WT secondary structure predicted via AlphaFold 2 are significantly different for mutations at WLYY sites (left) and sites 36 and 59, but not at sites 1 or 61.** Boxes show the interquartile range, and medians are indicated with horizontal lines within the boxes. $p$-values indicating statistically significant differences between distributions calculated according to the one-sided two-sample Wilcoxon rank-sum test.

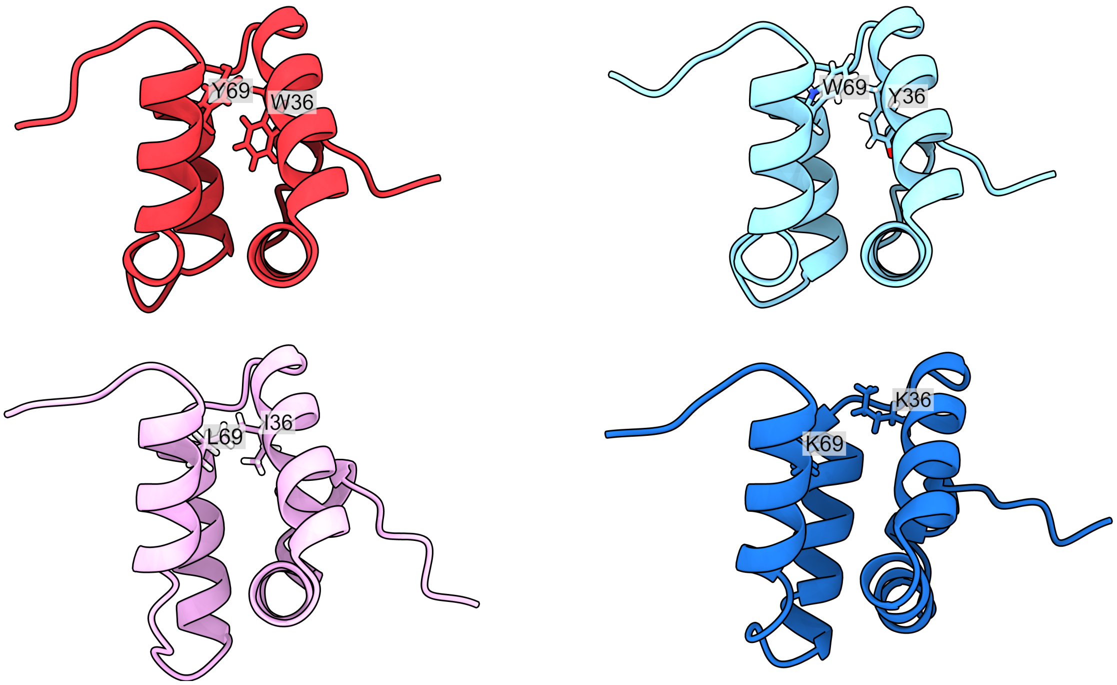


Figure S8: **AlphaFold 2 predictions for sequences with mutations at sites 36 and 69: 1) wild type - W at site 36, Y at site 69 (red), 2) mutant with Y at site 36, W at site 69 (cyan), 3) mutant with I at site 36, L at site 69 (pink) 4) mutant with K at sites 36 and 69 (blue).** Swapping the residues at sites 36 and 69 preserves the hydrophobic interactions that keep two helices together and results in a nearly unchanged tertiary structure (TM-align score vs. the WT: 0.98). Changing both residues to other hydrophobic ones (36I, 69L) results in a structure very similar to the WT (TM-align score: 0.85). Replacing both residues with similarly charged hydrophilic ones perturbs the spatial arrangement of the helices and results in a structure that diverges much more strongly from the WT (TM-align score: 0.64). Residues 36 and 69 are shown explicitly to highlight the increased distance between them in the last mutant. Images created with UCSF ChimeraX version 1.10 [3].
